# Large-scale deep tissue voltage imaging with targeted illumination confocal microscopy

**DOI:** 10.1101/2023.07.21.548930

**Authors:** Sheng Xiao, William J. Cunningham, Krishnakanth Kondabolu, Eric Lowet, Maria V. Moya, Rebecca Mount, Cara Ravasio, Michael N. Economo, Xue Han, Jerome Mertz

**Affiliations:** Department of Biomedical Engineering, Boston University, Boston MA 02215; Neurophotonics Center, Boston University, Boston MA, 02215

## Abstract

Voltage imaging with cellular specificity has been made possible by the tremendous advances in genetically encoded voltage indicators (GEVIs). However, the kilohertz rates required for voltage imaging lead to weak signals. Moreover, out-of-focus fluorescence and tissue scattering produce background that both undermines signal-to-noise ratio (SNR) and induces crosstalk between cells, making reliable *in vivo* imaging in densely labeled tissue highly challenging. We describe a microscope that combines the distinct advantages of targeted illumination and confocal gating, while also maximizing signal detection efficiency. The resulting benefits in SNR and crosstalk reduction are quantified experimentally and theoretically. Our microscope provides a versatile solution for enabling high-fidelity *in vivo* voltage imaging at large scales and penetration depths, which we demonstrate across a wide range of imaging conditions and different GEVI classes.

## Introduction

Instruments capable of monitoring the activity of large numbers of neurons with genetic specificity are crucial for the study of brain function^1^. While traditional patch-clamp electrophysiology is generally regarded as the gold standard for high-fidelity membrane voltage measurement, it is difficult to perform in vivo and mostly limited to single cells only. High-density multi-electrode arrays have expanded this capacity to hundreds of cells, though without the possibility of genetic specificity^2^. To date, only optical approaches have enabled the monitoring of neuronal activity with genetic specificity^3^,^4^. High-performance genetically encoded calcium indicators have enabled routine recordings of intracellular calcium dynamics across thousands of neurons^1^. However, calcium activity is a surrogate for the more fundamental electrical activity of neurons^5^. To directly capture membrane potential, genetically encoded voltage indicators (GEVIs) are required^6^, whose development has been an area of intense and ongoing activity. The latest generation of GEVIs can easily resolve individual action potentials with millisecond precision^7^–15, with sensitivities adequate to monitor subthreshold membrane potential variations.

Despite the enormous potential of GEVIs, their application to *in vivo* voltage imaging remains fraught with challenges. When attempting high-speed imaging over large fields-of-view (FOVs), signal levels per pixel are inevitably small, making it difficult to maintain adequate signal-to-noise ratio (SNR). This difficulty is exacerbated by tissue scattering and out-of-focus fluorescence that lead to background contamination and signal crosstalk (i.e. spurious signal from other neurons). To address these challenges, enhancements in GEVI performance, together with soma targeting^16^ and/or sparse labeling^17^, have made it possible to simultaneously image tens of cells in vivo using widefield microscopes equipped with state-of-the-art high-speed cameras featuring near perfect quantum efficiencies (QEs). However widefield imaging fails to provide background rejection, making it vulnerable to crosstalk and reduced SNR in cases of denser labeling. Targeted illumination can help limit the generation of out-of-focus background^10,18,19^, but becomes less effective with increasing target density. Alternatively, background can be inherently rejected with laser scanning microscopy (LSM), such as confocal^20^ or two-photon^11,15,21–23^ microscopy (2PM). However, most LSMs suffer from limitations in single-pixel detector sensitivity (<40% QE) and laser scanner throughput, both of which undermine SNR and FOV. To date, the ability to routinely perform sustained large-scale *in vivo* voltage imaging in densely labeled tissues with high SNR remains highly challenging.

To meet this challenge, we developed a kilohertz-rate **t**argeted **i**llumination **co**nfocal (TICO) microscope that combines the distinct advantages of widefield and scanning microscopy while circumventing their drawbacks. Our system incorporates technical innovations that simultaneously enable high fluorescence detection efficiency, high degree of background rejection, low photobleaching rates, kilohertz frame rates, wide imaging FOVs, and large penetration depths. We experimentally quantify the advantages of our microscope across multiple performance metrics in live mouse brains, which we supplement with a general theoretical framework for its optimization for *in vivo* applications.

We demonstrate the versatility of TICO microscopy with both a fully genetically encoded sensor somArchon^9^ and a hybrid chemogenetic sensor Voltron2^14^, under a variety of imaging conditions and across multiple brain regions, with FOVs as large as 1.16 × 0.325 mm and imaging speeds up to 4 kHz. At relatively superficial layers (< 200 µm) we demonstrate sustained voltage imaging of over 50 neurons in densely labeled tissue over recording durations of 20 minutes. The imaging depth is further extended for more sparsely labeled tissue regions, where we demonstrate high SNR voltage imaging of cortical layer 3 neuronal populations. Still deeper imaging is demonstrated of cortical layers 1-5 neurons through an implanted microprism, and of hippocampal CA1 neurons through an imaging cannula. With the combined advantage of large FOV, high SNR and background rejection, TICO microscopy allowed not only the simultaneous observation of distinct spiking activities across multiple cortical layers, but also spatially varying subthreshold oscillations across large neuronal populations, while at the same time achieving the highest single-photon voltage imaging depth of 300 µm in the mouse brain.

## Results

### TICO microscope design

The idea of integrating targeted illumination and confocal gating into a single instrument is guided by the fact that, while both techniques are effective at reducing out-of-focus background, they operate on complementary components of the image formation process: targeted illumination suppresses background generation by limiting out-of-focus excitation; confocal gating rejects out-of-focus fluorescence in the detection path. When combined, the techniques operate in synergy, thus minimizing background and its associated shot noise and signal contamination [Fig. 1(a-d)]. This principle is validated by a theoretical model that incorporates both targeted illumination and confocal gating, enabling their respective contributions in improving SNR and signal-to-background ratio (SBR) to be separately quantified for *in vivo* voltage imaging (Supplementary Note 1.1,1.2, Fig. S1,S2). Our results confirm that the optimized combination of targeted illumination and confocal gating offers higher SNR and SBR than either strategy alone (Supplementary Note 1.3, Fig. S3).

**Figure 1.**
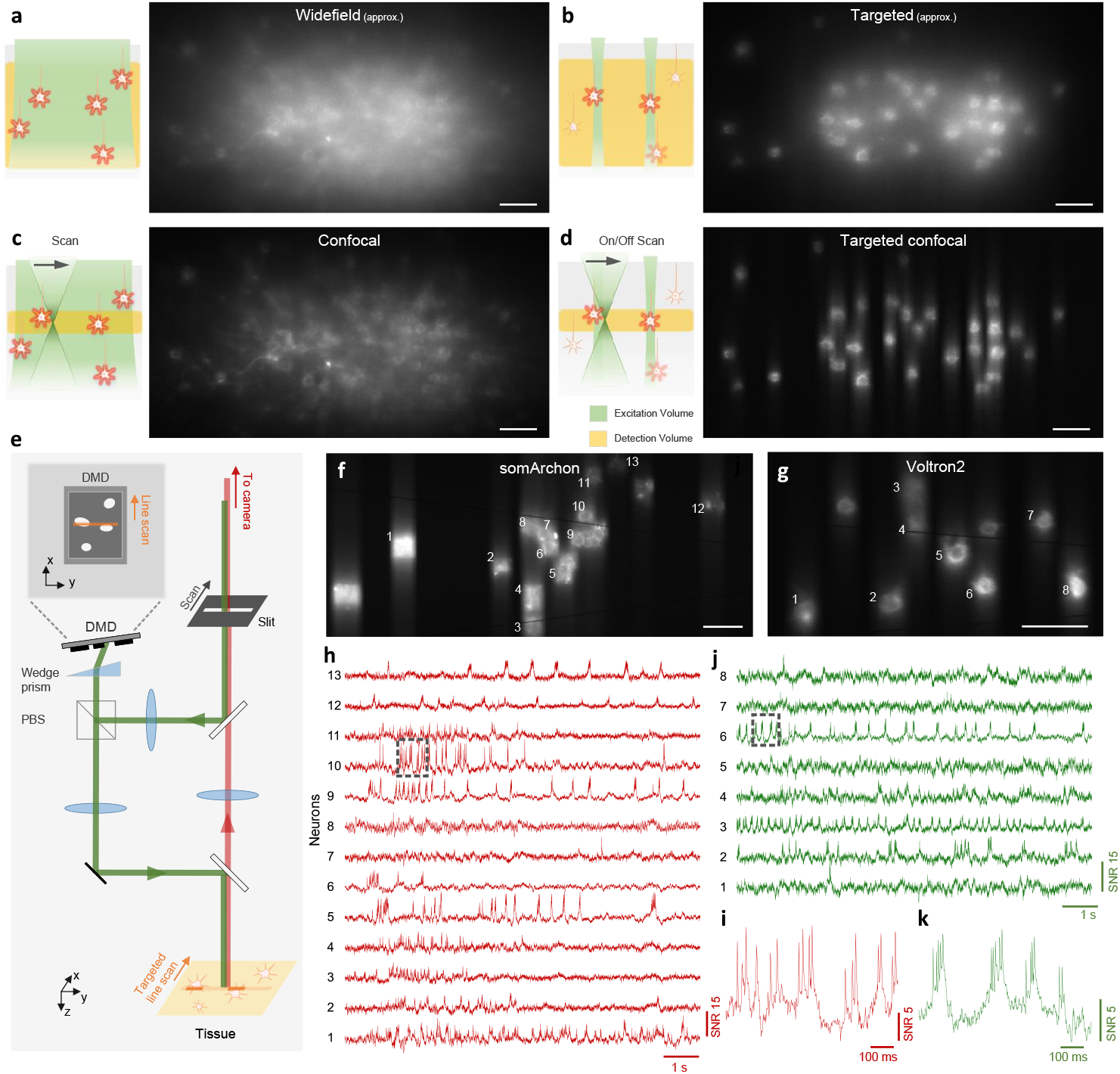
Principle and design of TICO microscope. (a-d) Illustration (left panel) and example images (right panel) of different excitation and detection strategies applied to *in vivo* voltage imaging of Voltron2: (a) standard widefield microscopy where all neurons in a volume are uniformly illuminated and detected with a widefield camera; (b) targeted illumination microscopy where only selected neurons within the focal plane are illuminated, but fluorescence from the entire volume is detected; (c) confocal microscopy where all cells within the volume are illuminated but only fluorescence near the focal plane is detected; (d) TICO microscopy where only selected cells within the focal plane are illuminated, and only the generated fluorescence near the focal plane is detected. Note that images in (a,b) were acquired using a confocal slit size of 156 µm, which produces higher contrast than a true widefield microscope (see Fig. S4). (e) Schematic of TICO microscope. Note that in actual setup, slit is fixed and scanning is performed with galvanometers. PBS, polarizing beamsplitter. (f-k) Demonstration of *in vivo* voltage imaging of somArchon (f,h,i) and Voltron2 (g,j,k) in the neocortex using TICO microscopy. (f,g) Averaged fluorescence images from somArchon and Voltron2. (h,j) Fluorescence traces from the active neurons labeled in (f) and (g) respectively. (i,k) Zoomed-in fluorescence traces from the boxed regions in (h) and (j) respectively. Scale bars in (a-d, f, g) are 50 µm.

To achieve confocal imaging over large scales and at high speeds, we implemented a line-scan strategy^24^. This not only enables millimeter FOVs to be sampled at kilohertz rates by 1D scanning, but also allows the use of large-aperture scanners to achieve high numerical aperture fluorescence collection. However, a drawback of most LSMs is their low detection efficiency that comes from the use of single-pixel detectors or a line-scan camera. We addressed this issue by additionally implementing a re-scan strategy where the fluorescence is re-imaged onto an area-scan camera by a second galvanometric scanner, thus enabling us to benefit from the exceptionally high QE of modern sCMOS cameras. Still, line-scan illumination alone does not maximize the efficiency with which excitation power is delivered to the sample, and line confocal gating alone does not maximize the suppression of background. To achieve these, we complemented our microscope with targeted illumination.

To enable targeted illumination, we inserted a digital micromirror device (DMD) in an intermediate image plane in the illumination path^19^. The illumination beam thus scans over a user-defined target pattern imprinted on the DMD, which is then projected into the sample with high contrast. While in conventional confocal microscopy the fluorescence detection path retraces the illumination path to achieve synchronized scan/de-scan^24^, here such a path overlap would severely undermine SNR because of diffraction losses introduced by the DMD [typically ∼40-50% ; Supplementary Note 3, Fig. S8, Fig. S9(a)]. In our own microscope, the minimization of fluorescence loss was a critical design consideration. We therefore decoupled the illumination and detection paths [see Fig. S9(b)], enabling the fluorescence signal to be de-scanned without passing through the DMD, resulting in a loss of only 6% due to the addition of two dichromatic mirrors. However, a technical complication arising from this strategy is that the illumination and detection planes are no longer co-planar in the sample, leading to a mismatch, both lateral and axial, of the illumination and detection FOVs that prevents effective confocal gating. While the lateral FOV mismatch can be corrected by placing the DMD in a Littrow configuration^25^, the axial mismatch is more difficult to correct, with approaches typically involving the addition of a matched grating^26^ or a multi-pass geometry^27^. Our own solution is simpler and more light efficient, and corrects the DMD-induced image plane tilt with a single wedge prism, which is capable of tilting the image plane by an angle *θ*_*w*_ = *δ* (*n*^2^ − 1)*/n* with an apex angle *δ* and refractive index *n*^28^ [illustrated in Fig. S9(c)]. When the DMD is tilted by this same angle, the excitation line remains in focus as it is swept across the DMD surface, allowing high-resolution light patterning across the entire FOV (Fig. S10). After beam reflection from the DMD, the backward path through the same wedge prism cancels this tilt such that the image plane becomes perpendicular again with respect to the propagation direction, restoring full confocality between the excitation and detection beams.

Following the design principles described above, we built a TICO microscope with decoupled illumination and detection paths using a DMD and off-the-shelf wedge prism [Fig. 1(e), Fig. S9(d)]. We confirmed the system’s confocality and capacity to target/image with cellular resolution over a FOV of 1.16 × 0.325 mm (Fig. S11). Three excitation wavelengths (488 nm, 561 nm and 637 nm) were integrated to enable multicolor imaging of either somArchon-GFP or Voltron2-JF552 [Fig. 1(f-k)]. Added flexibility was provided by the use of an adjustable confocal slit (0 - 156 µm width), which allowed us to readily control the degree of confocal sectioning. As shown below, this flexibility in both illumination targeting and confocal sectioning was critical in allowing the system to adapt to different sample conditions for maximum SNR.

### Characterization of TICO microscopy for *in vivo* voltage imaging

The purpose of combining targeted illumination with confocal gating is to increase the fidelity and duration with which neuronal activity can be monitored and quantified. By imaging Voltron2-expressing neurons using the same laser intensity at the brain surface but different microscope configurations (different confocal slit widths, with or without targeted illumination), we experimentally quantified the advantages of TICO microscopy over standard targeted illumination and confocal microscopy for *in vivo* voltage imaging. As is apparent from Figs. 1(a-d) and 2(a-d), both the application of targeted illumination and the strength of confocal gating affect spatial image contrast. When combined in TICO microscopy these provide a more than 50 × SBR improvement over conventional widefield microscopy (Supplementary Note 2.1, Fig. S4, S5). This capacity for background reduction was also reflected in a higher temporal signal contrast, as characterized by ∆*F/F* associated with individual spikes. With targeted illumination and moderate confocal gating (10 - 20 µm slit width) we were able to obtain an average spike ∆*F/F* as high as 8 - 10 % even in densely labeled tissue [Fig. 2(e,f)], which has only been reported previously with background-free neuron cultures^14^. We found TICO microscopy was able to significantly reduce background-induced crosstalk caused by both tissue scattering and out-of-focus neurons (detailed in Supplementary Note 2.2, Fig. S6), as is evident from Fig. 2(l-n).

**Figure 2.**
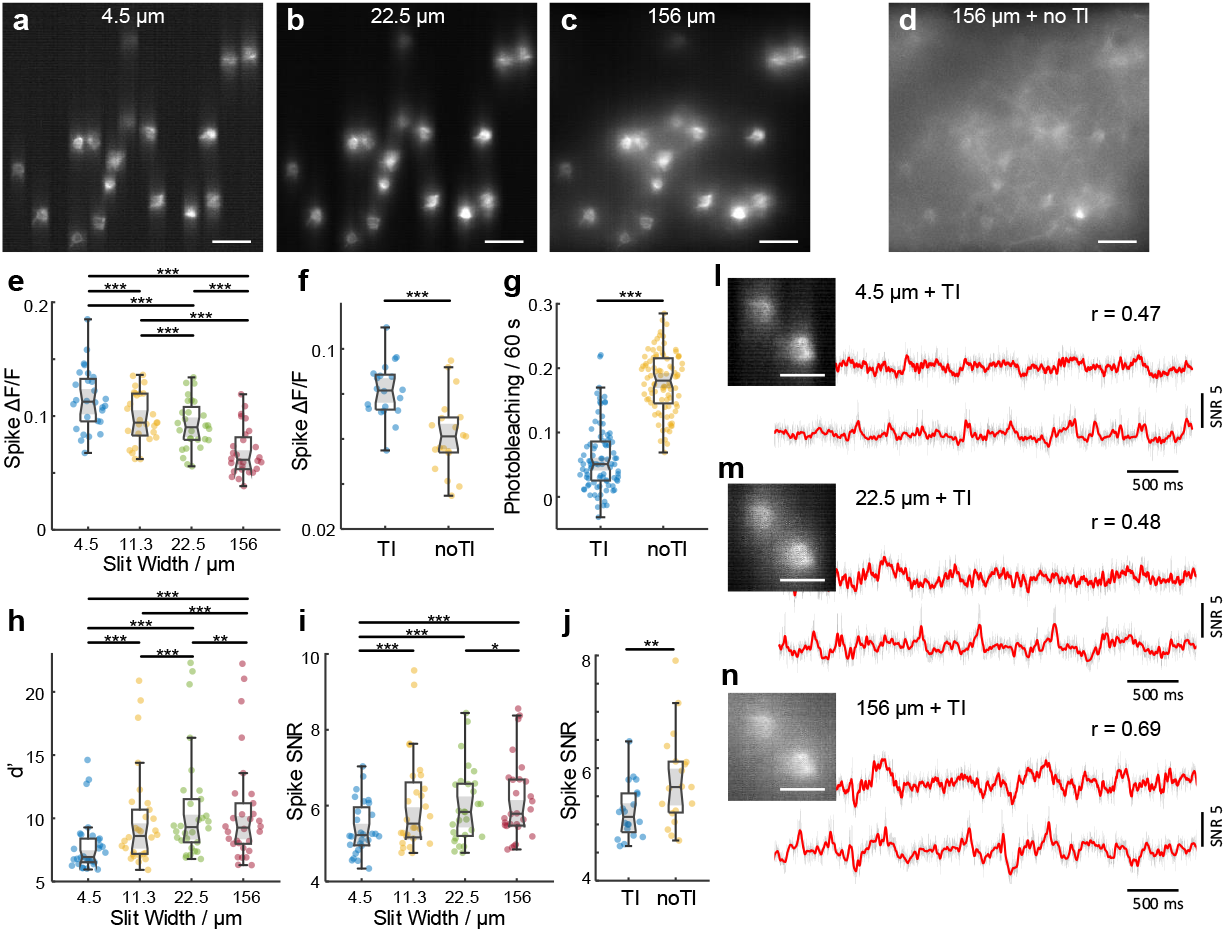
Quantification of TICO microscope performance for *in vivo* voltage imaging. (a-c) Example Voltron2 fluorescence images under targeted illumination with confocal slit width set to 4.5, 22.5, and 156 µm. Scale bar 50 µm. (d) Voltron2 fluorescence image over the same FOV but acquired without targeted illumination and with a confocal slit width of 156 µm. TI, targeted illumination. Scale bar 50 µm. (e,h,i) Comparison of spike ∆*F/F*, spike detection fidelity *d*^′^, and spike SNR measured with targeted illumination and confocal slit widths of 4.5, 11.3, 22.5, and 156 µm (n = 30 cells from 6 FOVs, 2 mice). Box plots: box, 25th (Q1, bottom line) to 75th (Q3, top line) percentiles; whiskers, *Q*1 − 1 5 × *IQR* to *Q*3 +1 5 ×*IQR*, where *IQR = Q*3 − *Q*1; middle line, median (m); notch, from 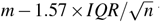 to 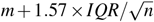; dots, measurement points. *p <* 0.05, *p <* 0.01, *p <* 0.001, no label if *p* 0.05, pairwise Wilcoxon signed-rank test, see Table S4 for statistics. (f,g,j) Comparison of spike ∆*F/F*, photobleaching rate, and spike SNR measured with and without targeted illumination when using a 14 µm confocal slit. For (f,j), n = 19 cells from 5 FOVs, 2 mice. For (g), n = 92 cells from 5 FOVs, 2 mice. (l,m,n) Example images (scale bar, 20 µm) and corresponding fluorescence traces from two neighboring neurons with targeted illumination and confocal slit widths of 4.5, 22.5, and 156 µm (from top to bottom). Gray line, fluorescence traces; red line, extracted subthreshold Vm traces; r, Pearson cross-correlation coefficient between the subthreshold Vm traces from the 2 neurons.

Another important factor for high-fidelity recording is SNR, which we quantified using both the theoretical shot-noise-limited spike detection efficiency *d*^′^ (see Supplementary Note 4 and Ref.^29^) and the experimental spike SNR (defined as spike amplitude over baseline noise, see Methods). As with conventional confocal microscopy^24^, an optimization of the confocal gating strength is required to achieve optimal SNR, which was attained in our case with a slit width of about 20 µm [Fig. 2(h,i)]. However, this optimum was found to be only weakly peaked and tolerant to a relatively wide range of slit widths, permitting a preference for a slightly stronger confocal gating for reduced crosstalk (Supplementary Note 1.3.1, 2.3). When targeted illumination was applied in addition, we observed a large reduction in photobleaching rate of 71.4% [Fig. 2(g)], caused by the reduction in received excitation power by the targeted neurons [Fig. S1(f)]. Despite this large reduction in excitation power (and hence fluorescence), the spike SNR was found to degrade only slightly from 5.66 to 5.13 [Fig. 2(j)], owing to the increased signal collection efficiency and background rejection resulting from the addition of targeted illumination [Supplementary Note 2.3, Fig. S7]. That is, TICO microscopy provides a capacity for long-duration imaging by virtue of significantly reduced photobleaching rates that in our case more than compensated the observed small degradation in SNR compared to confocal microscopy alone.

### Large-scale voltage imaging with different GEVI classes

State-of-the-art GEVIs can be either fully genetically encoded or hybrid, and differ in brightness, photostability, kinetics, voltage sensitivity, and even signal polarity. As shown above, TICO microscopy delivers minimum amounts of excitation power while maintaining high signal contrast and SNR, ensuring compatibility with different GEVI types. Together with the advantage of large FOV, TICO microscopy enables routine large-scale voltage imaging that can be sustained over long durations.

To demonstrate this, we first imaged Voltron2-expressing neurons in cortical layer 2 of awake mice. We selected a confocal slit size of about 14 µm to balance SNR and crosstalk. Because of this larger slit size, residual background remained visible under confocal microscopy but was largely removed with the addition of targeted illumination [Fig. 3(a), Fig. S12(a)]. With 40 mW/mm^2^ excitation intensity at the brain surface, we were able to image 57 neurons over a FOV of 1.1 × 0.325 mm^2^ continuously for 20 minutes (Fig. 3, Fig. S12). Individual spikes and Vm depolarizations can be observed throughout the full length of the recording [Fig. 3(d-f)]. The total photobleaching was measured to be 0.34/0.25-0.40 [median/Q1-Q3; Fig. 3(h)] across all imaged neurons, leading to a downward trend in spike SNR from 6.32/5.86-6.80 to 5.24/4.83-5.67 [median/Q1-Q3; Fig. 3(g)]. A similar such demonstration was performed using the fully genetically encoded sensor somArchon, both in superficial layers of cortex and in the hippocampal CA1 region (Fig. S15, S16), where we were able to image more than 50 neurons. Benefiting from the large FOV, up to 78 neurons could be imaged simultaneously (Fig. S14, S13), demonstrating the ability of TICO microscopy to perform sustained large-scale voltage imaging even in densely labeled tissues.

**Figure 3.**
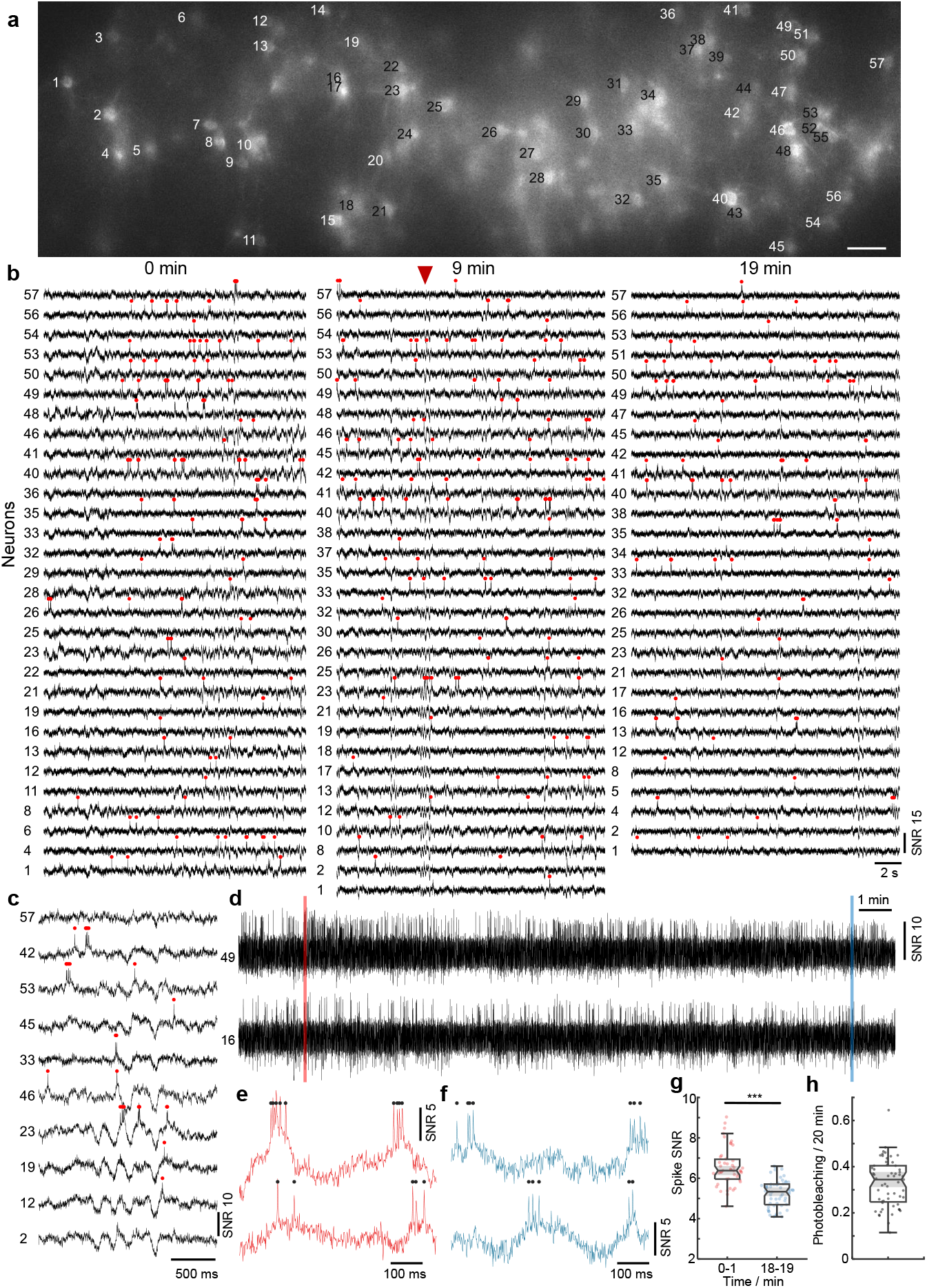
TICO microscope enables *in vivo* voltage imaging at large scales over extended durations. (a) Confocal image of Voltron2 fluorescence over the imaging FOV. Imaging depth 160 ± 20 µm, excitation intensity 40 mW/mm^2^, scale bar 50 µm. See Fig. S12(a) for the averaged Voltron2 fluorescence image over the same FOV with both confocal and targeted illumination. (b) Voltron2 fluorescence traces of active cells over 20 s periods starting at 0, 9, and 19 min in the recording. See Fig. S12(b) for the entire 20 min recording of all 57 cells. (c) Fluorescence traces from selected cells showing partially synchronized subthreshold oscillations. Extracted at the time point indicated by the red arrow in (b). (d) Example fluorescence traces from 2 selected cells during the entire 20 min recording. See Fig. S12(c) for raw fluorescence traces. (e,f) Zoomed-in fluorescence traces from the shaded areas labeled in (c). (g) Comparison of spike SNR during 0-1 and 18-19 min of the recording. ∗ ∗ ∗ *p* = 3.13*e*^−10^, Wilcoxon rank sum test, see Table S4 for statistics. (h) Measured photobleaching rate across all 57 imaged cells over the 20 min recording. Box plot same as Fig. 2(e). Median/Q1-Q3, 0.34/0.25 - 0.40.

Crucial to voltage imaging is the ability to detect subthreshold membrane potential oscillations. With TICO microscopy we were able to observe these among populations of neurons with high SNR using both Voltron2 and somArchon [Fig. 3(c), Fig. S17]. The advantages of high imaging throughput and low crosstalk were further highlighted by the observation of different oscillation patterns across distinct neuronal groups [Fig. 3(c)], as well as individual neurons that did not participate in this coordinated behavior [Fig. S17(c,h)]. Analysis from one of our animals showed a 3 -5 Hz central oscillation frequency, an association with hyperpolarization and reduced spike rates, and phase-locked spike timing to the oscillation cycles (Fig. S18). Similar 3 - 5 Hz membrane oscillations have previously been observed from neurons in cortical layer 2-6^30^–32, though mostly restricted to intracellular recordings of single neurons. Here, the high amount of temporal coordination of L1 membrane oscillations is in line with the proposed origin of thalamic axonal innervations^33,34^, although further study is required to elucidate its actual mechanisms.

### Deep tissue voltage imaging

Microscopes based on single-photon excitation provide only limited depth penetration in thick tissue because of scattering and out-of-focus background. In the case of voltage imaging, reported depth penetrations so far have been limited to typically around 100 - 150 µm^10,12,18,19,35^. Here, with the combined benefits of targeted illumination and confocal gating, we found that it is possible to extend this penetration depth to 300 µm, providing access even to cortical layer 3. The key challenge here stems from the increased light scattering that comes with increased imaging depth, rendering both background reduction mechanisms less effective. Regarding targeted illumination, tissue scattering blurs the excitation patterns such that they become less confined to the target neurons. As a result, reduced excitation power is incident on the targets [Fig. 4(n), Fig. S1(f)], which we compensate for by increasing the excitation intensity up to 80 - 150 mW/mm^2^ when imaging deep in tissue. Tissue scattering also blurs the emission signals, prescribing a larger confocal slit size to maintain optimized SNR, as predicted by our theoretical model [Fig. S2(h)]. In practice, we adopted a larger slit size of 23 µm when imaging deep in tissue. Because of the resultant increase in overall background fluorescence compared to in-focus signal, we intentionally sought to image more sparsely labeled brain regions where the SBR was naturally higher: the median SBRs across two typical FOVs at depths of 160 µm and 300 µm was 0.055 and 0.073 respectively when imaged without targeted illumination with a 156 µm confocal slit (Fig. S5). Note that for these experiments we did not specifically perform sparse labeling but rather used the same mice from the previous section, where Voltron2-expressing neurons tended to become more sparsely distributed as the imaging FOV moved away from the viral injection sites and approached cortical layer 4. Ultimately, we were able to routinely image populations of neurons at depths exceeding 200 µm (Fig. 4, Fig. S19; n = 12 FOVs from 3 mice), and up to 300 µm [Fig. 4(g,h,i), Fig. S19(c,d)]. Across all of our trials, the measured spike SNR and ∆*F/F* were 6.02/5.48-6.87 and 9.93%/7.61% - 11.76% (median/Q1-Q3; 61 active neurons out of 138 total neurons, 12 FOVs), comparable to our imaging results from more superficial layers.

**Figure 4.**
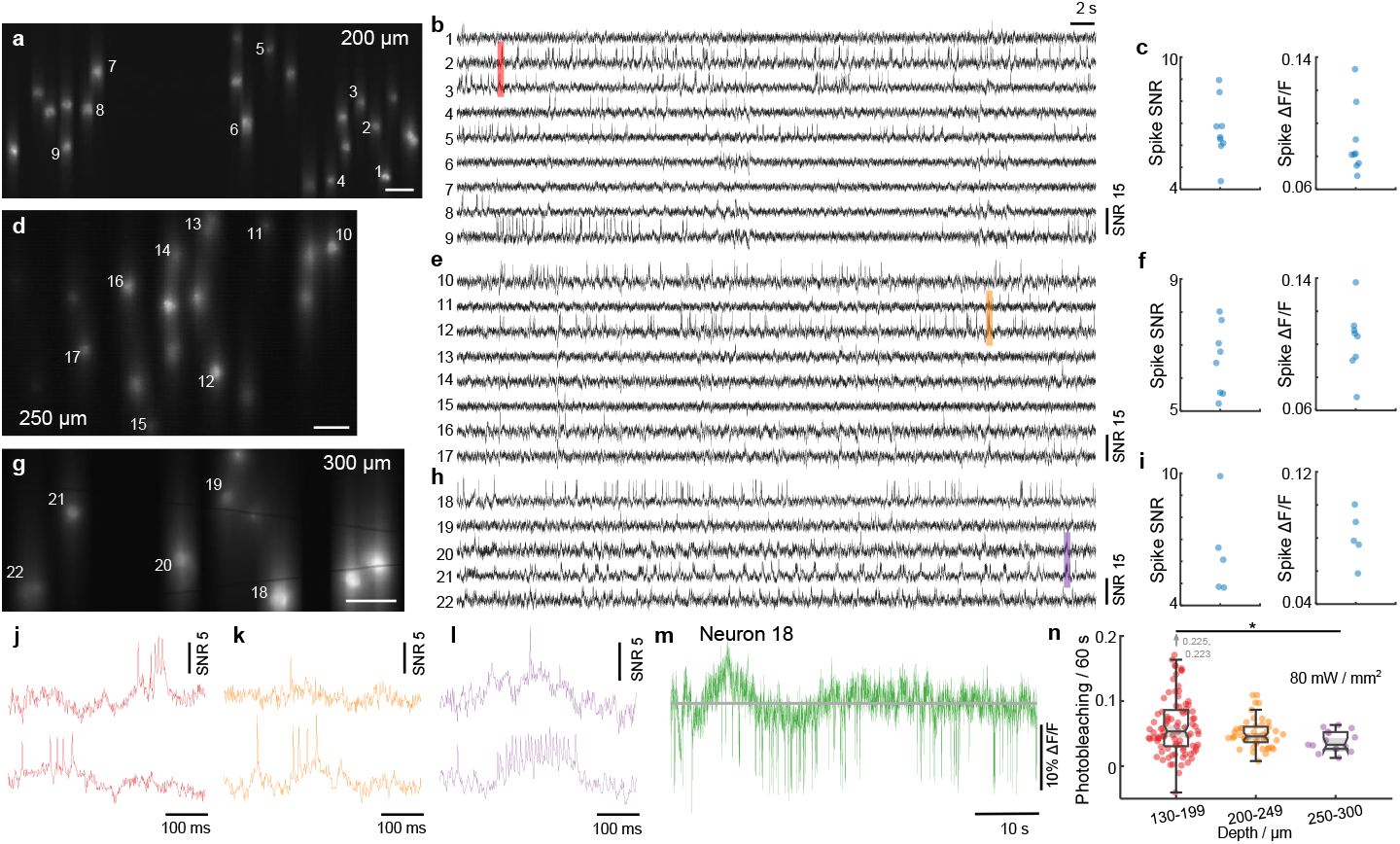
TICO microscope enables high SNR voltage imaging at 300 µm depth. (a-i) Example Voltron2 fluorescence images (a,d,g), voltage traces of spiking neurons (b,e,h), and average spike SNR, ∆*F/F* (c,f,i) at increasing imaging depths of 200 µm (a-c), 250 µm (d-f), and 300 µm (g-i) below the brain surface. Median spike SNR and ∆*F/F* are 6.37/0.081, 6.63/0.105, 6.08/0.078 for the active neurons shown in (b,e,h), respectively. See Table S3 for a list of imaging parameters. Scale bars in (a,d,g) are 50 µm. (j-l) Zoomed-in fluorescence traces from the shaded regions in (b,e,h). (m) Raw fluorescence trace of neuron 18 over the 1 min recording. (n) Comparison of photobleaching rate measured at different imaging depths. Excitation power density at the brain surface was kept constant at 80 mW/mm^2^. The reduction in photobleaching rate with increasing imaging depth is caused by the reduction in excitation power received by the targeted neurons due to tissue scattering. Box plot same as Fig. 2(e). ∗ *p <* 0.05, Wilcoxon rank sum test, see Table S4 for statistics.

### Simultaneous imaging across multiple cortical layers

Many deep brain regions such as the dorsal striatum and hippocampus are located well below the penetration depth limit of single-or even multi-photon microscopes^36^. To access these regions, a widely adopted strategy involves introducing tissue penetrating imaging conduits such as cannula^37^, gradient-index lenses^38^ or microprisms^39^. Our TICO microscope is entirely compatible with the use of such conduits, providing cellular resolution voltage imaging in densely labeled tissue in the hippocampus CA1 region (Fig. S16) and deep cortical layer 5 (Fig. S20).

A unique advantage of using an implanted microprism conduit is that it provides a side-on view of the brain. When coupled with a large imaging FOV, neural activity from an extended depth range can be recorded simultaneously. This is particularly important in brain regions such as neocortex, which is organized into multiple layers of distinct cell types and connectivity^40^. We injected Voltron2 virus at around 200 and 600 µm depth in the somatosensory mouse cortex, and implanted a right-angle prism of 1 × 1 mm^2^ facet size. Using TICO microscopy, we could simultaneously image over a vertical FOV of ∼ 800 µm spanning cortical layers 1 to 5 [Fig. 5(b)], almost the entire cortex. Distinct firing patterns were observed at different layers: while neurons in layer 2/3 (neuron # 2-20) mostly produced single, isolated spikes, 3 out of the 4 layer 5 neurons (neuron # 22-24) tended to produce more bursting events with substantial after-depolarizations. We note that owing to geometric constraints, the fluorescence collection efficiency was somewhat compromised when imaging away from the center of the microprism facet. Nevertheless, clear spike and subthreshold activity remained apparent even toward the top and bottom of the FOV [Fig. 5(d-f), Fig. S20].

**Figure 5.**
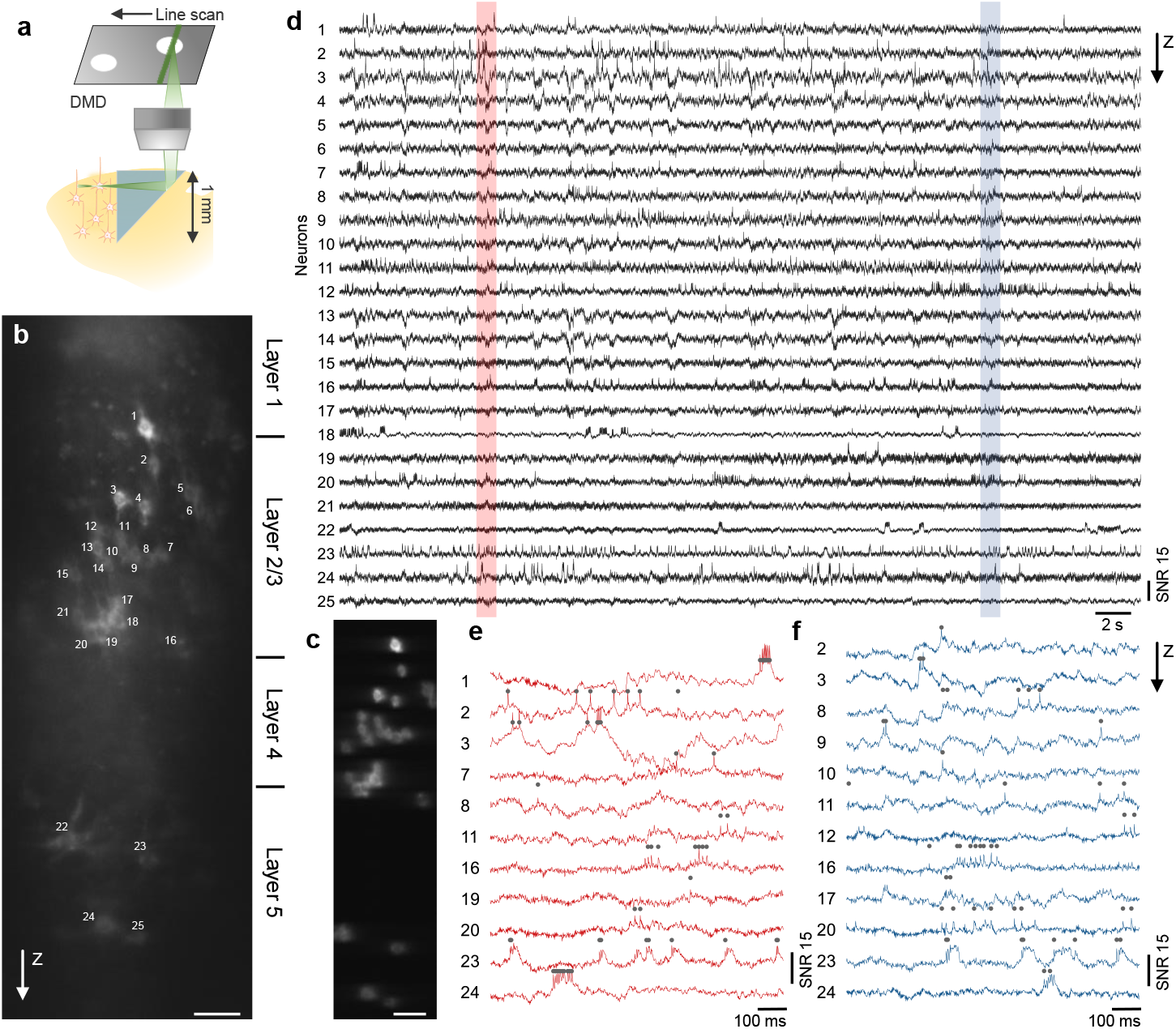
Side-on voltage imaging across multiple cortical layers with an implanted microprism. (a) Schematic illustration of side-on cortical imaging with an implanted microprism. (b) Side-on confocal image of Voltron2 fluorescence, demonstrating simultaneous view of neurons across cortical layers 1-5. Scale bar, 50 µm. (c) Averaged Voltron2 fluorescence image with 25 neurons targeted within the FOV. Scale bar, 50 µm. (d) Voltron2 fluorescence traces for the 25 targeted neurons shown in (b) over a continuous 45 s recording. (e,f) Zoomed-in fluorescence traces of active neurons in the shaded regions shown in (d).

## Discussion

TICO microscopy significantly improves upon targeted illumination or confocal microscopy in many aspects. Compared to confocal microscopy alone^20^, we were able to increase both fluorescence detection efficiency and imaging FOV by more than an order of magnitude using a combination of system designs including decoupled excitation/detection paths, a wedge prism for DMD tilt correction, and an adaptive confocal slit size that increases with deeper imaging. The addition of targeted illumination effectively compensated the weaker confocal gating that came with larger slit sizes, while also reducing the delivered excitation power and thus reducing photobleaching, enabling better compatibility with fully genetically encoded sensors. Compared to targeted illumination microscopy alone^10,14,19,^ a key advantage of adding confocal gating is to reduce crosstalk, enabling higher fidelity voltage imaging of large numbers of neurons. Improved SBR also facilitates the initial selection of in-focus neurons that can be difficult to distinguish from background when using widefield microscopy. As a point of clarification, when comparing the effects of confocal gating strength, we were unable to remove the gating completely due to a limitation in the maximum confocal slit size of 156 µm. And yet, as shown in Fig. S4, even this open slit led to at least 5.1× increase in SBR compared to a fully widefield microscope, allowing finer structures like proximal dendrites to become more easily identifiable. This suggests that the gains in SBR and SNR reported here are likely larger when compared to a fully widefield (no slit) targeted illumination microscope.

Another important metric when evaluating microscope performance is depth penetration, where 2PM is the most popular tool for deep imaging in scattering tissue. However, 2PM voltage imaging is limited by the requirement of exponentially increasing excitation power with depth and the moderate voltage sensitivity of currently available GEVIs, and to date has enabled only a few neurons to be imaged simultaneously^15,21,22^ at depths greater than 200 µm. In contrast, TICO microscopy is based on linear excitation which requires much lower laser power and benefits from the availability of better performing GEVIs, allowing such depths to be accessed routinely. A drawback is increased crosstalk at larger penetration depths due to tissue scattering, which can be alleviated by imaging more sparsely labeled tissue regions (Fig. 4), or, in the case of densely labeled tissue, by making use of imaging conduits, allowing voltage imaging at depths of several millimeters (Fig. S16, S20). Our results show that TICO microscopy, when applied to voltage imaging, can attain depths roughly on par with 2PM (even without the use of imaging conduits), while providing access to larger FOVs and better GEVI performance.

In our *in vivo* experiments we mostly set the confocal slit size to be between 11 - 23 µm, since for our demonstrations this provided near-optimal SNR with lower optical crosstalk. However, our system is highly flexible because it features an adjustable slit and DMD, meaning that both the strength of confocal gating and the fill factor of targeted illumination can be adapted to specific imaging tasks. For example, if one wishes to prioritize low optical crosstalk, the slit size can be narrowed to reduce background fluorescence. Alternatively, in the event neurons are sparsely labeled in layer 1 only, one can use an open slit with high-fill-factor targeted illumination for higher fluorescence collection efficiency.

Different from widefield-based microscopes^19^, the FOV and frame rate of our TICO microscope was limited here by the speed of our galvanometric scanners. We mostly focused on demonstrating large FOV imaging with 800 Hz acquisition frame rates, since this was sufficient for identifying individual spikes. This rate could be pushed to 1000 Hz over a FOV of 880 × 325 µm (Fig. S21). For applications where the exact determination of spike timing or spike width is important, faster imaging speeds of 2 kHz and 4 kHz were also achieved, though requiring reductions in FOV to 400 × 325 µm and 60 × 180 µm respectively (Fig. S22). Still faster frame rates for large FOVs could be envisioned with the use of resonant or polygonal scanners. Alternatively, large FOVs are more readily attainable when ultrafast imaging is not required. For example, TICO microscopy could be applied to other fluorescent sensors such as jGCaMP8^41^ or iGluSnFR^42^, which, while fast, are not as fast as GEVIs and can be imaged at reduced imaging speeds of about 100 Hz.

In summary, TICO microscopy is a practical and versatile solution for single-photon fluorescence imaging that can be adapted to a wide range of samples and imaging conditions. It provides the combination of low crosstalk and high SNR, while at the same time allowing large imaging FOVs, kilohertz acquisition speeds, low photobleaching rates, and large penetration depths, making it particularly suitable for general large-scale *in vivo* voltage imaging applications.

## Methods

### Imaging setup

#### TICO microscope

A detailed schematic of our TICO microscope setup is shown in Fig. S9(d), with all components listed in Table S2. In brief, the system was based on a line scan confocal microscope, where a line focus was imaged into the sample by way a series of relay lenses (*f*_6,7,8,9_) and an objective (Nikon 16×/0.8NA LWD). The line focus was scanned across the FOV by a galvanometric scanner (Galvo 1, ScannerMAX Saturn-5) placed in a plane conjugate to the objective back aperture. The generated fluorescence was epi-collected by the same objective, descanned by the same galvanometric scanner, and focused onto a stationary slit (Thorlabs VA100) that rejects out-of-focus fluorescence. The slit was then re-imaged onto an sCMOS camera (Teledyne Photometric Kinetix) by a pair of relay lenses (*f*_1,2_), and a second galvanometric scanner (Galvo 2, ScannerMAX Saturn-5) placed at the pupil plane re-scans the slit to form a 2D image on the camera. A combination of excitation filter, emission filter and dichromatic mirrors (Chroma Technology Corp., 89901v2 405/488/561/640nm Laser Quad Band Set) were used to separate fluorescence from the excitation light.

Different from a conventional line-scan confocal microscope is the addition of a DMD-based targeted illumination module, which was inserted between the scan lens *f*_4_ and the objective. To maximize collection efficiency, we avoided de-scanning the fluorescence through the DMD by separating the emission from the excitation light paths using two dichromatic mirrors (Chroma Technology ZT405/488/561/640rpcv2). That is, the DMD (Vialux V-7000 VIS) was present only in the excitation path to pattern the illumination beam. Specifically in the excitation path, the line focus was imaged and scanned across the DMD surface by a series of relay lenses (*f*_4,6,7,9_) and a galvanometric scanner (Galvo 1), which was then imaged into the sample by a tube lens *f*_8_ and the objective. The DMD was placed in the Littrow configuration and its surface tilt was corrected by a wedge prism (Edmund Optics # 49-443) inserted before it. A polarizing beamsplitter (PBS, Thorlabs WPBS254-VIS) and quarter-wave plate (*λ/*2, Thorlabs AQWP10M-580) were used to separate incident and reflected light from the DMD. An additional emission filter (Em2, Chroma Technology ZET405/488/561/640mv2) was placed in the emission path to block unpatterned excitation light transmitted through DM2.

We configured our microscope to feature three different excitation lasers. A 561 nm laser (Oxxius LCX-561L-200-CBS-PPA; 200 mW output power) was used for Voltron2 imaging. Its output was shaped into a line by a Powell Lens (Laserline Optics Canada, LOCP-8.9R10-1.0) and a cylindrical lens (*Cyl*_5_). Three additional cylindrical lenses (*Cyl*_5,6,7_) were used to further narrow the laser line focus. A 637 nm diode laser bar (Ushio America Inc., Red-HP-63X; 20 output beams, 6 W total output power) was used for somArchon imaging. Its output was collimated by a built-in fast- and slow-axis collimating lens, and two cylindrical lenses (*Cyl*_4,5_) were added to shape the output into a focused line. A 488 nm laser (Lasertack GmbH PD-01376) was used for GFP imaging, and three cylindrical lenses (*Cyl*_1,2,3_) were used to create a line focus. The outputs of the three lasers were combined using two dichromatic mirrors (DM4, Thorlabs DMSP550R; DM5, Thorlabs DMLP605R), and coupled into the microscope using a quadband dichromatic mirror (DM1, Chroma Technology Corp. ZT405/488/561/640rpcv2). Three half-wave plates (*λ*_1,2,3_*/*2) were placed at the output of each laser to rotate their beam polarizations such that the beams were reflected by the PBS towards the DMD in the excitation path.

#### Custom widefield microscope

A custom-built widefield epi-fluorescence microscope was used to compare spatial image contrast between standard widefield imaging and TICO microscopy with the largest slit width (156 µm) and without targeting. A 565 nm LED (Thorlabs M565L3) was collimated (Thorlabs ACL25416U-A), bandpass filtered (Chroma Technology Corp., ET560/40), and focused into the sample by way of an objective (Nikon 16×/0.8NA LWD). The generated fluorescence was collected by the same objective, separated from the excitation light using a dichromatic mirror (Chroma Technology Corp. T590lpxr), long-pass filtered (Chroma Technology Corp. ET590lp) and finally focused onto a sCMOS camera (Hamamatsu ORCA-Flash 4.0 V3) with a tube lens. Image acquisition was performed using HCImageLive software.

### Animal surgery

All animal procedures and experiments were carried out with approval from the Boston University Institutional Animal Care and Use Committee and in accordance with National Institutes of Health policies and guidelines. C57BL/6J mice (Jackson Laboratory #000664), CAG-Sun1/sfGFP mice (Jackson Laboratory #021039), and NDNF-ires-Cre mice (Jackson Laboratory #030757), both male and female, were used in this study.

#### Voltron2 mouse with crystal skull window

Both C57BL/6J and CAG-Sun1/sfGFP were used for Voltron2 cortical imaging. To allow optical access to the brain while minimizing damage, we followed methods similar to those described in Ref.^43^ for crystal skull preparation. Specifically, anesthesia was induced using 5% isoflurane in O_2_ and was maintained during surgical procedures using 1-2% isoflurane in O_2_. Bupivacaine (0.1 ml, 0.5%) was injected under the skin covering the skull. The skin and periosteum covering the skull were removed and the skull removed overlying the sites of interest. Virus was injected using a manual volume displacement injector (Narishige International USA, MMO-220A) connected to a glass pipette (Drummond Scientific, 5-000-2005) pulled to a 30 µm tip (Sutter Instrument, P-2000) that was beveled to a sharp tip. Pipettes were backfilled with mineral oil and virus was front-loaded before injection. Pipettes were inserted to the appropriate depth after the skull was removed.

Animals were unilaterally injected with a 1:12 dilution of AAV1-hSyn-FLEX-Voltron2-ST-WPRE virus in primary motor and primary somatosensory areas (coordinates in mm from Bregma: AP -0.5, ML +/-1.2, DV -0.3). Cortical injections to adult C57BL/6J mice additionally included a 1:300 dilution of rAAVretro-hSyn-Cre (Addgene #105553-AAVrg) to induce expression of Voltron2-ST and a 1:150 dilution of AAV9-hSyn-Cheriff-EGFP (Addgene plasmid #51697, custom viral preparation from The Penn Vector Core). Total injection volume at each target site was 50 nL. Following virus injection, the exposed area was then covered with modified crystal skull cover glass (LabMaker) and sealed with dental acrylic. To facilitate head fixation during imaging, we utilized a custom ring-shaped titanium head bar that was attached to the remaining cranial bone using low-viscosity cyanoacrylate adhesive (Loctite 4014) and dental acrylic. Mice were administered post-operative sub-cutaneous injections of ketoprofen (5 mg/kg) and buprenorphine (0.1 mg/kg) in saline for pain management. Viruses were allowed to express for 3 weeks before imaging was performed.

#### Voltron2 mouse with implanted microprism

C57BL/6J mice were used for Voltron2 cortical imaging via an implanted microprism. The surgical procedure to implant the glass microprism into the cortex was performed similarly to cranial window implant protocol detailed in Ref.^44^. Briefly, the animal was anesthetized and given pre-operative cefazolin and buprenorphine for pain management. The depilated scalp was resected to expose the entirety of the dorsal skull. A headbar was attached to the skull approximately over the lambda suture using cyanoacrylate glue (Loctite 4014) and dental acrylic, and the skin margins were secured to the outer edges of the skull using glue. A ∼ 3.5 mm diameter craniotomy was made over somatosensory cortex to accommodate the microprism assembly. A small lip was carved into the skull to allow the assembly cover glass to sit flush with the skull surface while resting on a thin layer of bone. The assembly was made by gluing a 1.0 mm glass microprism (hypotenuse coated with enhanced aluminum; Tower Optical MPCH-1.0) to a 3.5 mm diameter round coverglass using optical glue (Norland Products NOA61). Before implanting the assembly, 100 nL of AAV1-hSyn-FLEX-Voltron2-ST-WPRE mixed with 1:50 retroAAV-hSyn-Cre was injected into 3 sites across the craniotomy, at approximately 200 µm and 600 µm depths at each injection site. Virus was allowed to settle for approximately 15 minutes after each injection. A straight-line incision into the cortex was made using a micro scalpel blade mounted to the stereotaxic manipulator. The scalpel was inserted into the cortex first to a depth of 200 µm, and then translated in a straight line across ∼ 1.0 mm of cortex. The scalpel was removed and then reinserted at the start position of the incision to a depth of 400 µm and a 1.0 mm incision was made again. This process was repeated at 750 µm and 1000 µm. The assembly was then positioned by slowly pressing the microprism edge into the incised cortex until the cover glass sat flush with the skull surface. A plastic pipette tip was attached to the stereotaxic manipulator and used to maintain downward pressure on the assembly while the brain tissue settled around the implanted prism. The edges of the cover glass were secured to the skull using cyanoacrylate glue, and the entire skull cap was then covered with dental acrylic while ensuring that the cover glass remained exposed. Animals were given post-operative buprenorphine and ketoprofen for 2 days following the surgery and carefully monitored for 3 days after the procedure to ensure full recovery. Imaging was performed after 4 weeks following implantation.

#### SomArchon mouse with cortical window

NDNF-ires-Cre mice were used for somArchon cortical imaging. The surgical procedure for implanting an imaging window and a head-plate was detailed in Ref.^9^. The imaging window consisted of a circular coverslip (#0, outer diameter 3mm, Deckgläser Cover Glasses, Warner Instruments 64-0726). Virus injection and imaging window placement were performed under 1-3% isoflurane anesthesia, with sustained buprenorphine administered preoperatively to provide continued analgesia for 72 hours (buprenorphine hydrochloride, 0.03 mg/kg, i.m.; Reckitt Benckiser Healthcare). A craniotomy ∼ 3 mm in diameter was made near the right visual cortex. AAV virus was then infused via a blunt 36-gauge stainless steel needle (World Precision Instruments NF36BL-2) connected to a microliter injection system (10 µL, World Precision Instruments), attached to a stereotactic holding arm and controlled by a microinjector pump (World Precision Instruments UltraMicroPump). AAV virus was injected in 6 different locations across the craniotomy. The needle terminated about 250 µm below the dura. 300 nL of AAV9-Syn-FLEX-SomArchon-GFP (titer: 1.28 × 10^13^ GC/mL) was diluted 1:10 using a sterile saline solution and then infused at a rate of 50 nL/min, and the infusion cannula was left in place for 5 min at the end of the infusion to facilitate AAV spread. The imaging glass window was then positioned in the craniotomy, and surgical silicone adhesive (World Precision Instruments Kwik-Sil) was applied around the edges of the imaging window to hold it in place. Dental cement (Stoelting Co.) was then gently applied to affix the imaging window and a custom aluminum headbar posterior to the imaging window.

#### SomArchon mouse with implanted hippocampal cannula

C57BL/6J mice were used for somArchon hippocampal imaging. Animals underwent one surgery including stereotaxic viral injection targeting the hippocampus and implantation of a sterilized custom imaging cannula (outer diameter: 3.17 mm, inner diameter: 2.36 mm, height: 2 mm). The imaging cannula was fitted with a circular coverslip (size 0, outer diameter: 3 mm; Warner Instruments D263), adhered to the bottom using a UV-curable optical adhesive (Norland Products NOA60). During surgery, an approximately 3 mm circle was outlined on the skull (centered at anterior/posterior: -2.0mm, medial/lateral: +1.8mm). Three injections of 200 nL AAV9-CaMKII-SomArchon-GFP virus, obtained from University of North Carolina Vector Core (titer 3.2 × 10^12^ GC/mL), were made within this circle. Injections were performed with a blunt 33-gauge stainless steel needle (World Precision Instruments NF33BL-2) and a 10 µL microinjection syringe (World Precision Instruments NanoFil), using a microinjector pump (World Precision Instruments UMP3 UltraMicroPump). The needle was lowered over 1 min and remained in place for 30 sec before infusion. The rate of infusion was 50 nL/min. After each infusion, the needle remained in place for 7 min before being withdrawn over 1 min. After injections, an approximately 3 mm craniotomy was created using the outlined circle created previously. The cortical tissue overlaying the hippocampus was aspirated away to expose the corpus callosum. The corpus callosum was then thinned until the underlying CA1 became visible. The imaging cannula was then tightly fit over the hippocampus and sealed in place using a surgical silicone adhesive (World Precision Instruments Kwik-Sil). The imaging window was secured in place using bone adhesive (C&B Metabond Parkell) and dental cement (Stoelting Co.). A custom aluminum headplate was also affixed to the skull anterior to the imaging window. Analgesic was provided for at least 48 hours after each surgery, and mice were single-housed after window implantation surgery to prevent damage to the headplate and imaging window.

#### *In vivo* imaging

All videos were acquired using Teledyne Photometrics PVCAM software and recorded in RAW format for postprocessing. During the acquisition, the camera was freely running at a preset frame rate in either “sensitivity” or “speed” mode. The output trigger from the camera was used to synchronize the galvanometer scanning with a multifunctional DAQ card (National Instrument USB-6343), which produced a smoothed triangular wave defined as *y* = *A* arcsin[(1 − *ξ*) sin(2*πt f*)] that controlled the line scan position (*A* is the scan amplitude, *ξ* = 0.03 is the smoothing factor, *f* is the camera frame rate, *t* is time). The DMD was controlled using a custom Matlab script based on Vialux ALP-4.2 API.

#### SomArchon mouse imaging

Before imaging, mice were head-fixed under the microscope objective while allowed to freely run on a floating Styrofoam ball. Because GFP and somArchon were co-localized at the soma, imaging areas were identified using GFP fluorescence under 488 nm laser excitation, ∼ 14 µm confocal slit size, and no targeted illumination. Based on the confocal GFP fluorescence image, small rectangular ROIs encompassing the cell bodies were manually drawn for the in-focus neurons. This created a binary mask that was uploaded to the DMD for excitation light patterning. SomArchon voltage imaging was performed using the 637 nm laser according to the parameters listed in Table S3.

#### Voltron2 mouse imaging

Prior to imaging, a retro-orbital injection was performed to deliver 100 µL of solution containing 20 µL of Pluronic F-127 (20% w/v; Sigma Aldrich), 20 µL of DMSO, and 100 nmol of Janelia Fluor 552 dye in sterile PBS.

Imaging was performed 1-3 days after the dye injection. During the imaging sessions, mice were awake, head-fixed under the microscope objective while constrained within a 1-inch acrylic tube. Before acquisition, imaging areas were identified using Voltron2 fluorescence with laser power attenuated using a reflective neutral density filter (Thorlabs ND20A, 1% transmission) to avoid photobleaching. For each imaging area, a confocal image acquired without targeted illumination was used as a reference to select illumination targets as described previously. After removing the neural density filter, Voltron2 voltage imaging was performed using the same 561 nm laser according to the parameters listed in Table S3.

For comparisons of imaging performance using different microscope configurations (Fig. 2), we imaged the same neurons under each configuration by interleaving 10 s long trials. For each FOV, each configuration was imaged for a total of 30 - 40 s over 3-4 individual trials. Imaging depths ranged from 130 to 200 µm. To estimate photobleaching in Fig. 2(g) and Fig. 4(n), the trials were extended to 60 s but only imaged once per FOV. This produced more photobleaching, allowing increased measurement accuracy. The excitation intensity at the brain surface was kept at 80 mW/mm^2^.

### Data analysis

#### Video preprocessing and voltage signal extraction

All recordings were saved in RAW format using Teledyne Photometrics PVCAM software. Each recording was first corrected for global motion using a masked object Fourier domain cross-correlation algorithm^45^, where the registration mask was manually selected based on the most distinguishable features within the averaged frame. From each video, the ROI of each neuron was manually selected, with voltage signals extracted by averaging all pixels values within the ROI at each time point. A camera offset was subtracted from the signal (20 ADU for “speed” mode and 100 ADU for “sensitivity” mode). An experimentally calibrated camera conversion gain (0.793 e-/ADU for “speed” mode and 0.264 e-/ADU for “sensitivity” mode) obtained using the mean-variance technique^46^ was used to convert ADU values into actual number of detected photons. During experiments we found that the peak QE of the camera was measured to be ∼ 95% under “sensitivity” readout mode, but only ∼ 45% under “speed” readout mode due to a design flaw in the readout electronics.

#### Spike detection and spike SNR estimation

Spike detection and SNR estimation were performed using the same procedure outlined in Ref.^9,19^. The algorithm consisted of several steps, including estimating baseline noise *σ*_*F*_ (*t*), spike detection and estimating subthreshold traces *F*_*sub*_(*t*). Since this algorithm assumes action potentials produce positive changes in fluorescence, all Voltron2 fluorescence traces were inverted before calculation.

From the raw fluorescence trace *F*_*raw*_(*t*), we first removed photobleaching by high pass filtering *F*_*raw*_(*t*) at 1 Hz, resulting in a detrended fluorescence trace *F*_*detrend*_(*t*). This was further highpass filtered at 50 Hz to remove subthreshold fluctuations, resulting in a trace containing both noise and potential spikes *F*_*hp*_(*t*). The baseline noise was estimated as twice the standard deviation of the downwardly rectified trace of *F*_*hp*_(*t*) over a local time window of ±1 s:

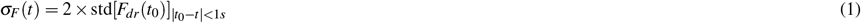

where 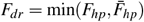 is the downwardly rectified trace, and 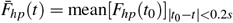 is locally averaged over a moving window of ± 0.2 s. Here by calculating the noise from only the downwardly rectified trace, we reduced bias in the noise estimation due to spike signals.

To find spike locations, we similarly generated an upwardly rectified trace containing all potential spikes: 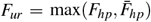. Since individual action potentials are characterized by sharp rises in fluorescence intensity, we calculated the temporal changes of upwardly/downwardly rectified traces as *dF*_*ur,dr*_(*t*) = *F*_*ur,dr*_(*t*) − *F*_*ur,dr*_(*t −* ∆*t*). Spike locations *t*_*AP*_ were determined from the time points that jointly satisfy:

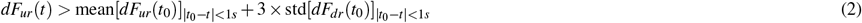

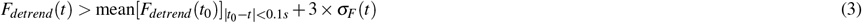

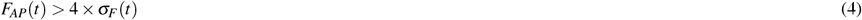

where *F*_*AP*_(*t*) is the spike amplitude, calculated as the maximum signal rise within 3 ms before the spike time:

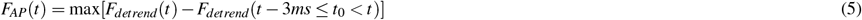

With the detected spike locations, we then produced spike-removed raw fluorescence traces *F*_*nospike*_(*t*) by replacing the intensities around the spike locations (1 ms before and 2 ms after the spike) with average the fluorescence intensities within a local ± 5 ms time window from the raw trace. Baseline fluorescence *F*_0_(*t*) was estimated by lowpass filtering *F*_*nospike*_(*t*) at 1 Hz, and subthreshold Vm traces *F*_*sub*_(*t*) were estimated by bandpass filtering *F*_*nospike*_(*t*) between 1 - 50 Hz. Throughout the manuscript ∆*F/F* traces were calculated as (*F*_*raw*_ *− F*_0_)*/F*_0_, SNR traces were calculated as *F*_*detrend*_*/σ*_*F*_, spike ∆*F*_*AP*_*/F* and SNR were estimated as *F*_*AP*_(*t*_*AP*_)*/F*_0_(*t*_*AP*_) and *F*_*AP*_(*t*_*AP*_)*/σ*_*F*_ (*t*_*AP*_).

To estimate photobleaching, we fitted the fluorescence baseline trace *F*_0_(*t*) to an exponential function *f* (*t*) = *a* · exp(*b* · *t*). The amount of photobleaching over a duration ∆*t* was then calculated as 1 − *f* (∆*t*)*/a*.

#### Estimation of spike detection fidelity

We calculated the theoretical shot-noise-limited spike detection fidelity *d*^′^ by adapting the framework developed in Ref.^29^ to the case of voltage imaging with a scanning microscope. This is detailed in Supplemental Text 4. Briefly, at each spike location *t*_*AP*_, we calculated the theoretical spike amplitude from the measured spike ∆*F/F* according to the relationship:

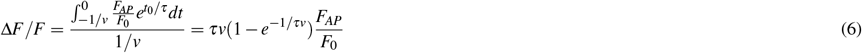

under the assumption of a fluorescence signal model of 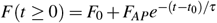, where *F*_0_ is the baseline fluorescence, *F*_*AP*_ is the spike amplitude, *t*_0_ [1*/v*, 0] is the spike onset time, *v* is the sampling rate, *τ* is the decay time of the fluorescent indicator (here assumed to be *τ* = 0.8 ms for Voltron2). Spike detection fidelity is then calculated as

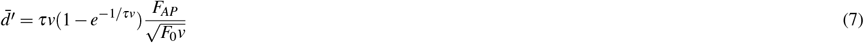

#### Estimation of optical crosstalk

To calculate ∆*F*_*r*_*/*∆*F*_0_ as a function of distance from the cell membrane (Fig. S6), we first gradually expanded the ROI of each neuron by performing morphological dilation (Matlab function *imdilate*) with a 7-pixel radius disk structural element, allowing us to obtain 7-pixel wide (3.16 µm) donut-shaped ROIs surrounding the same neuron with increasing distances from the cell membrane. At each spike time point *t*_*AP*_, ∆*F*_0_ was determined according to Eq. 5 within the central ROI, whereas ∆*F*_*r*_ was determined from surrounding donut ROIs at the same time points.

To analyze Vm-Vm correlations, we calculated Pearson cross-correlation coefficients (Matlab function *corrcoef*) for the extracted subthreshold traces *F*_*sub*_(*t*) from pairs of neurons. Their separation distances were calculated as the distance between centroids of respective ROIs.

#### Estimation of image contrast

We estimated the image contrast for each neuron as *SBR* = (*µ*_*s*_ − *µ*_*b*_)*/µ*_*b*_, where *µ*_*s*_ is the average intensity within the neuron ROI, and *µ*_*b*_ is the average intensity from a donut ROI surrounding the neuron. The donut ROI was obtained by taking the differences between the original ROI and a morphologically dilated ROI. To account for the anisotropic background distributions resulting from the use of a confocal slit, the morphological dilation was performed using an elliptical structural element with 5 µm vertical axis, and 2 µm horizontal axis.

#### Frequency-resolved analysis of subthreshold traces

To calculate Vm power at each frequency step *f*_0_, we applied a 2nd-order Butterworth filter with a lower and higher cutoff frequency of 0.8 *f*_0_ and 1.2 *f*_0_ to the spike-removed fluorescence trace *F*_*nospike*_(*t*). The analytical signal was then derived by Hilbert transformation to obtain phase and power.

To select of time periods of high/low Vm power in the network, we first averaged the Vm power across simultaneously recorded neurons to obtain a population-averaged Vm power trace. This was done based on the observation that the Vm delta oscillations were highly correlated across neurons. Normalized population Vm power in the delta frequency range was defined here to be between 2 - 5 Hz. Periods with Vm power below and above 2 standard deviations from the distribution were classified as low and high Vm power periods respectively.

#### Computation of spike-Vm phase locking

To quantify how consistent spikes occurred relative to the oscillation phase for each neuron, we first calculated the phase-locking value^47^ (PLV) defined as:

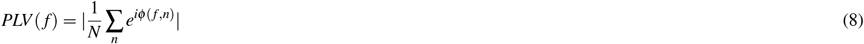

where *φ* (*f, n*) is the phase of the n-th spike at frequency *f* obtained by Hibert transformation, *n* = 1, …, *N*, and *N* is the total number of spikes. To further account for any potential differences in the number of spikes between groups of neurons, we adopted the unbiased phase locking value (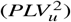) from Ref.^48^:

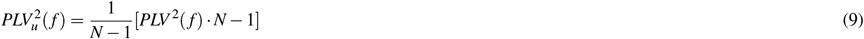

#### Statistical analysis

All statistical analysis was performed using Matlab 2021b. For comparison of spike amplitude, spike detection fidelity, spike SNR and spike ∆*F/F* in Fig. 2, Fig. S7, only neurons with measured spike rates of at least 1 Hz for at least one of the investigated imaging conditions (one of the slit widths or with/without targeted illumination) were included to minimize the effects of false-positiv
e spikes. For comparison of Vm-Vm correlation, ∆*F*_*r*_*/*∆*F*_0_, and photobleaching, all neurons within the FOV were included. The Wilcoxon signed-rank test was used for paired data (Matlab function *signrank*), and Wilcoxon rank sum test was used for unpaired data (Matlab function *ranksum*). The following applied for all box plots in the manuscript: box, 25th (Q1, bottom line) to 75th (Q3, top line) percentiles; whiskers, *Q*1 − 1 5 × *IQR* to *Q*3 + 1 5 × *IQR*, where the interquantile range *IQR* = *Q*3 − *Q*1; middle line, median (m); notch, from 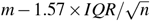 to 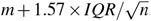; dots, measurement points or outliers according to the figure caption.

## Acknowledgements

This work was supported by NIH grants R34NS127098, R01MH122971, RF1MH126882, and F32MH129149.

## Author contributions statement

S.X., X.H. and J.M. conceived the project. S.X. designed and built the TICO microscope. W.J.C., K.K., M.V.M. and R.M. prepared experimental animals. S.X. performed imaging experiments with assistance from W.J.C., K.K., M.V.M, and C.R. S.X. and E.L. analyzed the data. S.X. wrote the manuscript with contributions from W.J.C., E.L. and R.M. M.N.E., X.H. and J.M. edited the manuscript. All authors reviewed the manuscript. M.N.E., X.H. and J.M. supervised the project.

## Data availability statement

Data underlying the results presented in this study will be made publicly available on Zenodo.

## Code availability statement

All relevant code for data processing will be available at https://github.com/HanLabBU.

## Competing interests

The authors declare no competing interests.

## Notes

### Competing Interest Statement

The authors have declared no competing interest.

