## Supplementary Material for "Large-scale deep tissue voltage imaging with targeted illumination confocal microscopy"

### Supplementary Information

#### 1 Theoretical study of fluorescence imaging using targeted illumination and confocal gating

##### 1.1 Theory

In this section we develop a theoretical model for fluorescence imaging with targeted illumination and confocal gating. We consider a generalized model of scanning microscopes for all imaging configurations, as shown in Fig. S1(a). In this model, an excitation focus is scanned in 2D across the sample, with or without targeted illumination, and fluorescence is detected through an adjustable confocal gate. We assume the sample has a fluorophore distribution described by $O(\vec{r})$, and the excitation and detection PSFs are described by normalized circularly symmetric $PSF_e(\vec{r})$ and $PSF_d(\vec{r})$, where $\int PSF_{e,d}(\vec{\rho},z)d\vec{\rho} = 1$, $\vec{r} = (\vec{\rho},z) = (x,y,z)$ is the 3D coordinate. During imaging, the excitation intensity distribution when the laser beam is scanned at location $\vec{\rho}_0$ can be written as

$$\begin{aligned} I_e(\vec{r}_s,\vec{\rho}_0) &= M_T(\vec{\rho}_s)\delta(\vec{\rho}_s-\vec{\rho}_0)\otimes PSF_e(\vec{r}_s) \\ &= M_T(\vec{\rho}_0)PSF_e(\vec{r}_s-\vec{\rho}_0) \end{aligned} \tag{S1}$$

where $M_T(\vec{\rho})$ is the 2D targeted illumination mask, $\vec{r}_0$ and $\vec{r}_s$ are spatial coordinates at the DMD and the sample plane, and $\otimes$ represents a convolution. The generated fluorescence distribution in the sample is obtained by multiplying Eq. S1 by the sample fluorophore distribution, leading to

$$I_f(\vec{r}_s,\vec{\rho}_0) = I_e(\vec{r}_s,\vec{\rho}_0)\cdot O(\vec{r}_s) \tag{S2}$$

From here, we distinguish two different detection strategies to reflect differences in implementations of scanning microscopy, namely without and with fluorescence re-scanning[1].

Most commonly laser scanning microscopy is implemented without re-scanning, where the fluorescence signal is detected by a single-pixel detector (for point scan) or a line camera (for line scan), and the image is formed by numerically assigning intensity readout values according to the scan location $\vec{\rho}_0$:

$$\begin{aligned} I_d(\vec{\rho}_0) &= \int d\vec{\rho}_c A_d(\vec{\rho}_c-\vec{\rho}_0)[I_f(\vec{r}_c,\vec{\rho}_0)\otimes PSF_d(\vec{r}_c)] \\ &= \iint d\vec{\rho}_c d\vec{r}_s A_d(\vec{\rho}_c-\vec{\rho}_0)I_e(\vec{r}_s,\vec{\rho}_0)O(\vec{r}_s)PSF_d(\vec{r}_c-\vec{r}_s) \\ &= M_T(\vec{\rho}_0)\,[O(\vec{\rho}_0)\otimes[PSF_e(\vec{r}_0)\cdot[A_d(\vec{\rho}_0)\otimes PSF_d(\vec{r}_0)]]] \end{aligned} \tag{S3}$$

where $A_d(\vec{\rho})$ represents the detection aperture (Table S1), $\vec{r}_c$ is the coordinate at an intermediate image space for confocal gating, and $\vec{r}_d$ are the coordinates in the final detection space.

Alternatively, if re-scanning is implemented, a second set of scanners is used to optically assign fluorescent photons onto a 2D multi-pixel detector with pixel size assumed to be infinitely small[1]:

$$\begin{aligned} I_d(\vec{\rho}_c) &= \int d\vec{\rho}_0[A_d(\vec{\rho}_c-\vec{\rho}_0)\cdot[I_f(\vec{r}_c,\vec{\rho}_0)\otimes PSF_d(\vec{r}_c)]] \\ &= \iint d\vec{\rho}_0 d\vec{r}_s A_d(\vec{\rho}_c-\vec{\rho}_0)I_e(\vec{r}_s,\vec{\rho}_0)O(\vec{r}_s)PSF_d(\vec{r}_c-\vec{r}_s) \\ &= \int d\vec{r}_s O(\vec{r}_s)PSF_d(\vec{r}_c-\vec{r}_s)\int d\vec{\rho}_0 A_d(\vec{\rho}_c-\vec{\rho}_0)M_T(\vec{\rho}_0)PSF_e(\vec{r}_s-\vec{\rho}_0) \end{aligned} \tag{S4}$$

**Table S1.** Detection apertures for different imaging configurations. $v_d$ is the radius of confocal pinhole or half-width of the confocal slit.

| Imaging configuration | $A_d(\vec{\rho})$ |
|---|---|
| Point scanning confocal | $\lvert\vec{\rho}\rvert < v_d$ |
| Line scanning confocal | $\lvert x\rvert < v_d$ |
| Widefield | 1 |

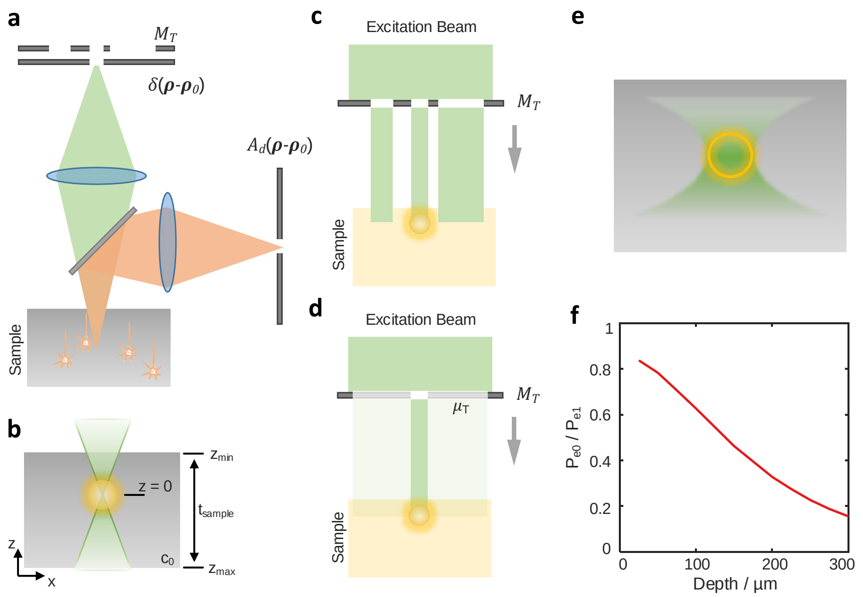


**Figure S1.** (a) Modeling for fluorescence imaging using targeted illumination and confocal gating. A targeted illumination mask $M_T$ is overlaid on top of the excitation mask $\delta(\vec{r}-\vec{r}_0)$ that controls the excitation pattern, with fluorescence signals spatially filtered by a detection aperture $A_d$.
(b) Imaging of fluorescence signal confined to cell membrane. Note that the fluorescent object is larger than the excitation focus.
(c) Patterning the excitation light using a binary targeted illumination mask.
(d) Patterning the excitation light using a grayscale targeted illumination mask. Central region of mask corresponding to the cell of interest has unit transmittance, with everywhere else having a transmittance equal to the average mask fill factor $\mu_T$.
(e) Illustration of reduced excitation power due to the use of targeted illumination. Because of tissue scattering and the finite depth-of-field of the microscope, the excitation power is reduced near the periphery of the cell. This is in contrast to non-targeted illumination where the entire cell receives the same amount of excitation power throughout.
(f) Reduction of excitation power under targeted illumination compared to non-targeted illumination at different imaging depths. Because scattering increases with depth, cells located deeper inside the tissue receive less excitation power.

Note that Eq. S4 can also be used for modeling a widefield microscope by setting the detection aperture $A_d(\vec{\rho}) = 1$. The models for these different imaging configurations are summarized in Table S1.

Here Eq. S3, S4 can be generally applied to different imaging configurations with varying degrees of confocal gating (by adjusting $A_d(\vec{\rho})$), with/without targeted illumination (by adjusting $M_T(\vec{\rho})$), and with/without image re-scan. Specifically, for standard confocal or widefield imaging without targeted illumination, we have $M_T(\vec{\rho}) = 1$ and therefore $I_e(\vec{r}_s, \vec{\rho}_0) = PSF_e(\vec{r}_s - \vec{\rho}_0)$. Thus Eq. S3, S4 can be reduced to

$$I(\vec{r}) = O(\vec{r}) \otimes PSF_{tot}(\vec{r}) \tag{S5}$$

where $PSF_{tot}(\vec{r}) = PSF_e(\vec{r}) \cdot [A_d(\vec{r}) \otimes PSF_d(\vec{r})]$ for confocal microscopy without re-scanning, and $PSF_{tot}(\vec{r}) = PSF_d(\vec{r}) \cdot [A_d(\vec{r}) \otimes PSF_e(\vec{r})]$ for confocal microscopy with re-scanning.

#### 1.2 Simulation details

We next aim to develop a simulation model relevant to *in vivo* voltage imaging conditions. To simulate soma-targeted membrane imaging [Fig. S1(b)], we assume the fluorescence signal can be modeled as a spherical shell of radius $r_{neuron} = 7.5$ µm, thickness $t_{neuron} = 4$ nm, and centered at $\vec{r} = (0,0,0)$:

$$O_s(\vec{r}) = \begin{cases} 1 & \text{if } r_{neuron} \le |\vec{r}| \le r_{neuron} + t_{neuron} \\ 0 & \text{elsewhere} \end{cases} \tag{S6}$$

In addition, we assume background is produced by a uniform fluorescent slab of finite axial span $z \in [z_{min}, z_{max}]$ and normalized fluorescence concentration $c_0 \in [0,1]$ outside the cell:

$$O_b(\vec{r}) = \begin{cases} 0 & \text{if } |\vec{r}| \le r_{neuron} \\ c_0 & \text{if } |\vec{r}| > r_{neuron} \text{ and } z \in [z_{min}, z_{max}] \end{cases} \tag{S7}$$

We estimate the background fluorescence concentration $c_0$ using anatomical data of the typical brain[2,3]. Specifically, we assume a cell density of 9.2e$^4$/mm$^2$, with each soma being a spherical shell of 15 µm diameter and membrane thickness of 4 nm. With perfect soma targeting and membrane localization, the fractional ratio of soma membrane within a unit volume is $1.1e^{-3}$. Furthermore, the fraction of cells labeled with GEVI can be affected by viral delivery and genetic targeting, which we assume to be $\mu_N \in [0.01, 1]$, leading to the normalized background fluorescence concentration $c_0 = 1.1e^{-3} \cdot \mu_N$. Bearing in mind that if we allowed the thickness of the background volume to be semi-infinite widefield microscopy would have infinite background and produce no contrast at all, we limited the background thickness to be $t_{sample} = z_{max} - z_{min} = 1$ mm, with the fluorescent object $O_s(\vec{r})$ located in the range 0 - 300 µm below the background volume surface $z = z_{min}$.

In the case of targeted illumination, typically a binary illumination mask is used that targets only in-focus objects. This leads to a spatially varying illumination pattern, and thus a spatially varying degree of background rejection. Here we adopt a simplified model to study the average effect of targeted illumination, where a gray-scale targeted illumination mask is defined as

$$M_T(\vec{\rho}) = \begin{cases} 1 & \text{if } |\vec{\rho}| \le r_{neuron} \\ \mu_T & \text{elsewhere} \end{cases} \tag{S8}$$

where $\mu_T \in [0,1]$ is the average fill factor of the targeted illumination mask. In Eq. S8 and illustrated in Fig. S1(c,d), the mask has unit transmittance within a central disk region corresponding to a targeted cell $O_s(\vec{r})$ of interest, allowing excitation light to fully reach the cell. Outside the cell, the mask has a reduced transmittance $\mu_T$ equal to the ratio of ON pixels to the total pixels of DMD (i.e., the fill factor), such that out-of-focus background from illumination targets other than $O_s(\vec{r})$ can be captured. Note that $M_T(\vec{\rho})$ can be further decomposed into a uniform mask $M_{T0}(\vec{r}) = 1$ (no targeted illumination) and targeted mask $M_{T1}(|\vec{\rho}| \le r_{neuron}) = 1$ (fully targeted illumination), where

$$M_T(\vec{r}) = \mu_T M_{T0}(\vec{r}) + (1-\mu_T) M_{T1}(\vec{r}) \tag{S9}$$

This allows us to further decompose the final detected fluorescence image into one generated by the uniform mask $M_{T0}(\vec{r})$ and targeted mask $M_{T1}(\vec{r})$:

$$I_s(\vec{r}) = \mu_T I_{s0}(\vec{r}) P_{e0} + (1-\mu_T) I_{s1}(\vec{r}) P_{e1} \tag{S10}$$

$$I_b(\vec{r}) = \mu_T I_{b0}(\vec{r}) P_{e0} + (1-\mu_T) I_{b1}(\vec{r}) P_{e1} \tag{S11}$$

where $I_s(\vec{r})$ is the signal image generated by $O_s(\vec{r})$, $I_b(\vec{r})$ is the background image generated by $O_b(\vec{r})$, and the subscripts $(*)_{0,1}$ on $I_{s,b}$ represent fluorescence images produced by the uniform and targeted masks respectively. Here we introduced a pair of new variables $P_{e0,e1}$ as the excitation power under non-targeted or targeted illumination, such that different amounts of ballistic excitation power can be delivered into the sample. If $P_{e0} = P_{e1}$, the cell may receive less excitation power when the illumination is targeted [Fig. S1(e,f)] due to a reduction of non-ballistic excitation. As a result, this can lead to perceived SNR differences, and manifested in our experiments as reduced photobleaching in the case of targeted illumination. By adjusting $P_{e0,e1}$ such that the cell of interest receives the same amount of excitation power under both non-targeted and targeted illumination:

$$P_{e0} \int d\vec{r} O_s(\vec{r}) = P_{e1} \int d\vec{r} [[M_{T1}(\vec{\rho}) \otimes PSF_e(\vec{r})] \cdot O_s(\vec{r})] \tag{S12}$$

we eliminate differences in photobleaching rate, allowing us to focus on the effects of fluorescence collection efficiency and background rejection on the final SNR.

For neuronal imaging, one is generally interested in the integrated signal within a predefined ROI (such as a soma). The ROI can be selected manually or with specialized algorithms. Here, with a knowledge of both signal and background, we always choose a circular ROI that maximizes the SNR in the final detected image $I_s(\vec{r}) + I_b(\vec{r})$:

$$SNR = |\alpha| F_s / \sqrt{F_s + F_b} \tag{S13}$$
$$F_s = \eta \int_{ROI} d\vec{r} I_s(\vec{r}) \tag{S14}$$
$$F_b = \eta \int_{ROI} d\vec{r} I_b(\vec{r}) \tag{S15}$$

where $\eta$ here is a scaling factor that converts recorded intensity into photon counts, and $\alpha$ is the percentage change of fluorescence signal from baseline.

Similarly, we can also study the amount of crosstalk induced by background fluorescence, which we define from the baseline-signal-to-background ratio:

$$SBR = F_s / F_b \tag{S16}$$

Here for voltage imaging we set $\alpha = 10\%$, and normalize the fluorescence signal with $\eta$ such the total emitted fluorescence from the cell membrane is $F_s = 10,000$. Therefore the theoretical maximum SNR is 10 for a widefield microscope with 100% detection efficiency and no crosstalk from background fluorescence ($SBR = +\infty$).

Finally, brain scattering plays an important part in *in vivo* imaging. Here we used NAOMi[4] to calculate the scattering-degraded PSFs for both excitation and detection at different depths inside the brain. This technique generates a simulated volume with refractive index variations based on actual anatomical data, including brain vasculature and random scatterers of varying size and strength. The 3D PSFs can then be obtained by numerically propagating the wavefront from the microscope back aperture through the simulated anatomical volume. We modified the original program to evaluate one-photon rather than two-photon PSFs, with all anatomical data kept as default. According to our experimental parameters, we approximated the excitation and detection wavelengths to both be $\lambda = 0.6$ µm, with excitation $NA_e = 0.4$, and detection $NA_d = 0.8$. The final scattering PSFs were averaged over 25 locations across the simulated volume. Tissue absorption was ignored.

##### 1.3 Simultation results

In this section we aim to provide a general guide for the optimization of TICO microscopy for *in vivo* imaging, and study how varying degrees of confocal gating $A_d(\vec{\rho})$ and targeted illumination $\mu_T \in [0.01, 1]$ affect the imaging performance in terms of SNR and SBR. We account for different imaging conditions by allowing for adjustments in imaging depth $z_{min}$ and labeling density $\mu_N \in [0.01, 1]$ (affecting both scattering and background fluorescence). To be in accord with our actual implementation of TICO microscopy, we confine ourselves here only to the re-scanned imaging model described by Eq. S4.

###### *1.3.1 Effects of confocal gating on SNR*

To maximize the SNR of a confocal microscope, the pinhole/slit size must be optimized to balance signal collection and background rejection[1]. In the case of a point-object model and in the absence of scattering[5], the optimal size of a confocal pinhole is found to match or be slightly larger than the excitation focus (1-3 Airy units for a diffraction-limited system). This principle is generally followed in most confocal imaging systems. However, conditions for *in vivo* imaging differ significantly from an ideal point-object model in that: (1) signal arises from an extended object (cell membrane in our case) that can have an axial extent larger than the microscope depth-of-field; (2) tissue scattering leads to blurred PSFs such that the system is no longer diffraction limited. Both of these factors suggest that to collect more signal one must increase the confocal pinhole/slit size beyond its conventional setting. However such an increase also leads to more background, bringing the overall effect on SNR into question.

To address this question, we start by investigating the effects of confocal gating on the attainable SNR under our simulated imaging conditions with no targeted illumination ($\mu_T = 1$) and a high labeling density $\mu_N = 1$. At each imaging depth, we calculate the SNR and SBR obtained from a cell of interest as a function of confocal pinhole/slit size $2v_d$. As with a standard confocal microscope, an increase in pinhole/slit size leads to an increase in background fluorescence as reflected by a decrease in SBR [Fig. S2(d-f)]. In terms of SNR, there still exists an optimal pinhole/slit size, albeit much larger than for the case of a point object: to achieve maximum SNR at 150 µm depth, the optimal $2v_d$ for point and line scan confocal are 19 and 16 µm respectively, instead of 1 and 0.8 µm in case of a point object embedded in a clear (non-scattering) medium. As the imaging

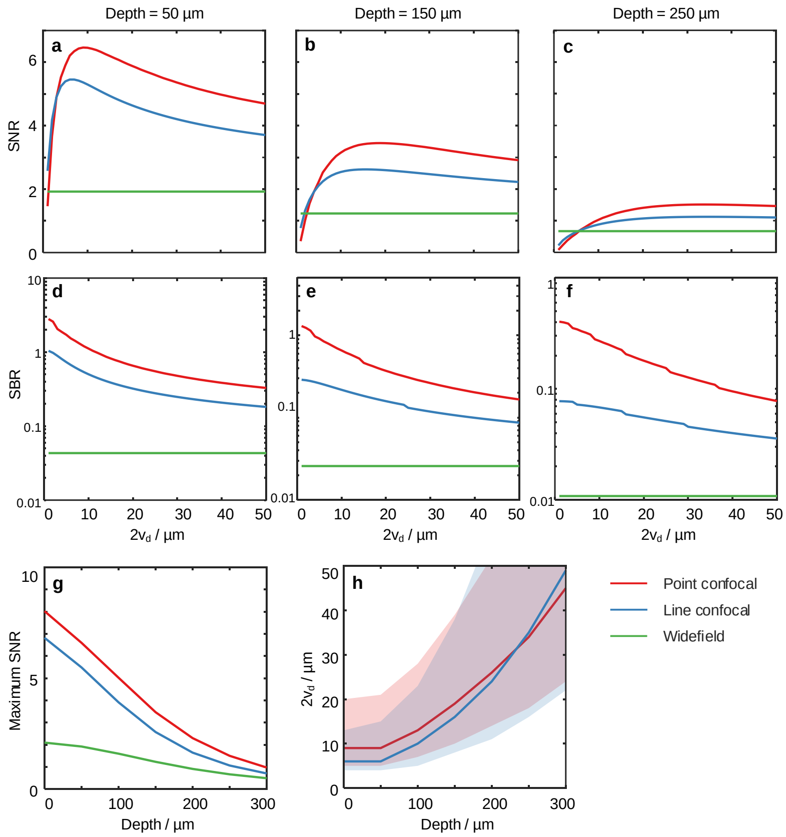


**Figure S2.** Effect of confocal gating strength on SNR. Simulation performed without targeted illumination ($\mu_T = 1$) and with labeling density $\mu_N = 1$.
(a-f) Imaging SNR (top row) and SBR (middle row) as a function of confocal pinhole/slit size $2v_d$ at different imaging depths.
(g) Maximum achievable SNR as a function of imaging depth across 3 imaging configurations with optimized confocal pinhole/slit size.
(f) Solid line, pinhole/slit size $2v_d$ to attain maximum SNR at varying depths. Shaded area, pinhole/slit size $2v_d$ to attain at least 90% of the maximum SNR.

depth increases, the confocal pinhole/slit must be opened further to accommodate the PSF blurring caused by tissue scattering [Fig. S2(h)], while the maximum achievable SNR decreases [Fig. S2(g)]. Note that at larger imaging depths, the SNR becomes only weakly dependent on $v_d$ once it reaches the shoulder in the curve, as shown in the shaded areas of Fig. S2(h) that represent all $2v_d$ values that attain 90% of the maximum SNR. That is,slightly smaller $v_d$ values can be used to reduce crosstalk with only minimal penalty on SNR.

##### *1.3.2 Optimal SNR and SBR with both confocal gating and targeted illumination*

Having established an approach to optimize the system SNR, we next seek to understand how the combination of targeted illumination and confocal gating influence SNR and crosstalk. Here for all imaging conditions, the confocal pinhole/slit size is optimized to achieve the maximum SNR. Figures S3(a-c) show the maximum SNR for the three considered imaging systems at 150 μm imaging depths with various tissue labeling densities and targeted illumination mask fill factors. In general, increasing imaging depth and labeling density all lead to reduced SNR, which can be alleviated with the application of either confocal gating or targeted illumination. While both techniques are effective in improving SNR, we found that the combined strategy shows marginal SNR improvements compared to fully targeted illumination if the excitation targets are sparsely distributed ($\mu_T = 0.01$). However, if the targeted illumination mask fill factor is increased to address a larger number of neurons, the addition of confocal gating provides a larger SNR benefit. In fact, with confocal gating, the SNR becomes almost invariant and close to optimal for targeted illumination masks with a moderate excitation density $\mu_T \leq 0.1$.

Another important consideration when evaluating single-photon imaging techniques is the crosstalk that arises from

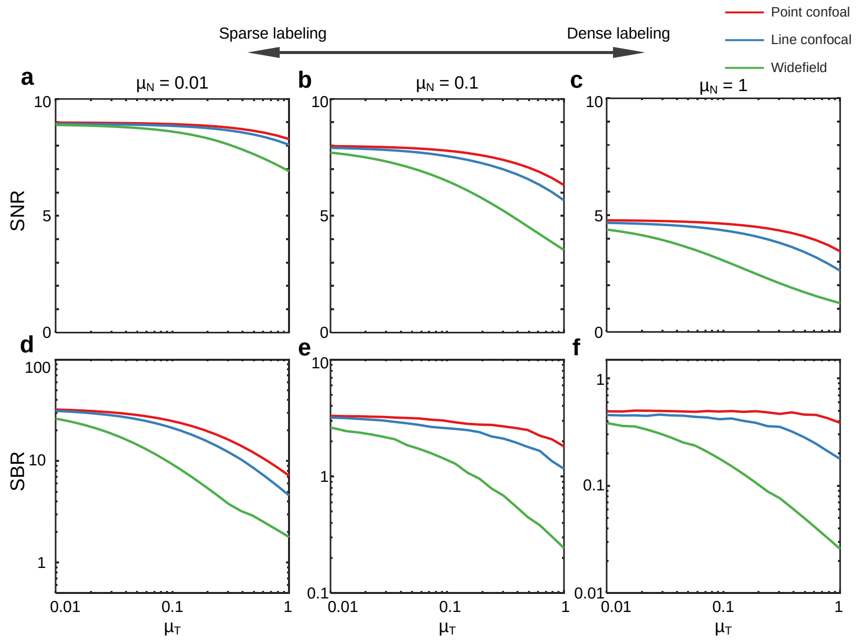


**Figure S3.** Comparison of theoretical SNR and SBR for *in vivo* voltage imaging with targeted illumination and confocal gating.
(a-c) Maximum SNR as a function of targeted illumination mask fill factor $\mu_T$ at 150 µm imaging depth with different sample labeling densities $\mu_N$.
(d-f) Theoretical SBR under the same conditions in (a-c) when maximum SNR is achieved.

background fluorescence, as characterized by SBR. At optimal SNR, SBR follows a similar trend where confocal gating is most beneficial when the excitation targets are densely distributed, but less so when the labeling density or targeting density is low [Fig. S3(d-f)]. However, with a small penalty on SNR, stronger background rejection can be achieved by using smaller confocal pinhole/slit sizes, which would lead to more significant SBR advantages even in case of sparse targeted illumination masks ($\mu_T = 0.01$).

### 2 Characterization of TICO microscopy for *in vivo* voltage imaging

#### 2.1 TICO microscopy improves image contrast

We begin by evaluating the respective benefits of targeted illumination and confocal gating on background reduction. These benefits are quantified most simply by their effect on the apparent image contrast as characterized by the SBR associated with cell bodies. We found that when performing *in vivo* imaging of Voltron2-expressing neurons at high labeling densities, individual neurons were barely distinguishable from background when using conventional widefield microscopy, even in regions where they were sparsely distributed [Fig. S4(a,d); median SBR 0.0151 for n = 61 neurons over 5 FOVs]. When confocal imaging was applied over the same FOVs, even a large slit size improved SBR considerably. We found that slit sizes of 156 µm and 11.3 µm (projected into sample) led to increases in SBR of 5.1× and 13.9× respectively (Fig. S4). This gain was further amplified 3.6× with the addition of targeted illumination [Fig. S5(a-d,i); n = 52 cells from 1 FOV, 14 µm confocal slit width], leading to overall improvements in SBR of $\sim 18\times$ and $\sim 50\times$, compared to conventional widefield microscopy. We note that this SBR improvement is likely an underestimate, since we were unable to identify individual neurons and locate the same FOV in more densely distributed regions with widefield microscopy, while we could routinely image these with TICO microscopy.

#### 2.2 TICO microscopy reduces crosstalk

While the SBR characterizes the spatial image contrast, more important for voltage imaging is the temporal fluorescence signal associated with individual neurons. A key requirement here for high-fidelity recording is that crosstalk between neurons be kept to a minimum. Two sources of crosstalk are: 1) fluorescence spread from nearby neurons due to tissue scattering, and 2) fluorescence from out-of-focus neurons, both of which lead to signal contamination. The observed reduction in background that comes from both targeted illumination and confocal gating is reflected in the increase in the temporal contrast $\Delta F/F$ of

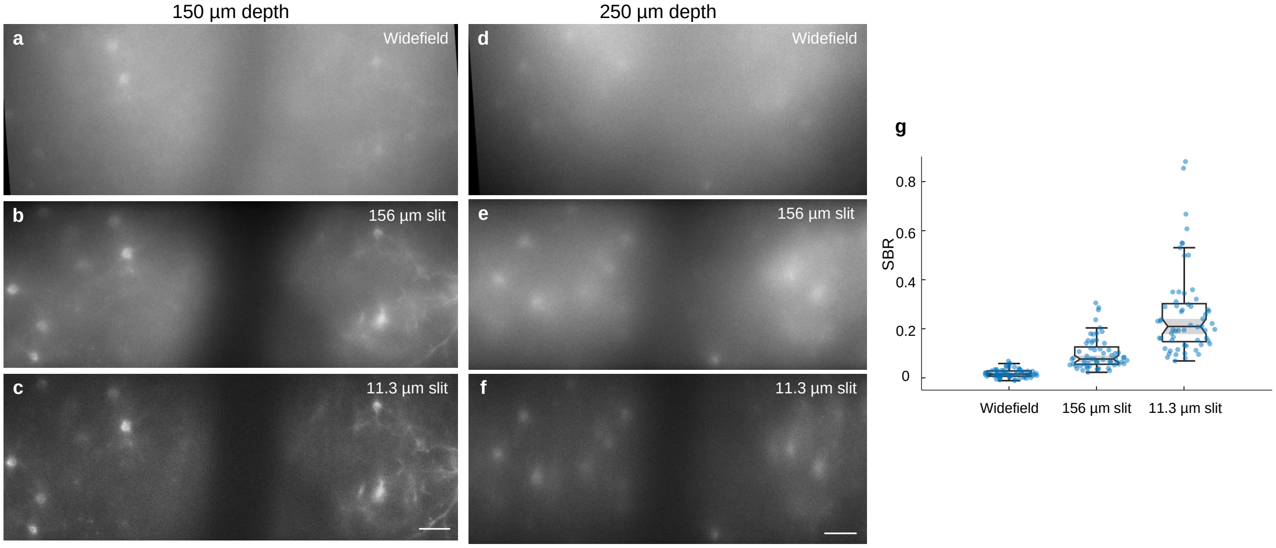


**Figure S4. Comparison of spatial image contrast under a standard widefield microscope and TICO microscope.**
(a) Voltron2 fluorescence image acquired with a standard widefield microscope using a LED for excitation and a sCMOS camera for detection.
(b) Voltron2 fluorescence image over the same FOV as (a), but acquired with TICO microscopy without targeted illumination and with a confocal slit size of 156 µm.
(c) Same as (b) but with a slit size of 11.3 µm. Scale bar 50 µm.
(d-f) Same as (a-c) but at an imaging depth of 250 µm.
(g) Comparison of SBR of neuronal somas under different microscope configurations. $n = 61$ cells over 5 FOVs at depths in the range 130 - 250 µm. The median/Q1-Q3 for widefield, 156 µm confocal slit and 11.3 µm confocal slit are 0.0151/0.0066-0.0283, 0.0768/0.0553-0.1259, and 0.2093/0.1474-0.3022. Box plot the same as in Fig. 2(e).

individual spikes [Fig. 2(e,f)], from which we expect a commensurate reduction in crosstalk. Here we evaluate the two sources separately in detail.

To quantify the spread of fluorescence from a neuron due to tissue scattering, we evaluated the decay of the measured spike amplitude as a function of distance from the neuron. Specifically, we measured $\Delta F_r/\Delta F_0$, where $\Delta F_0$ is the spike amplitude at the neuron location, and $\Delta F_r$ is the spike amplitude away from the neuron, averaged over annular ROIs of increasing radius. In the absence of scattering where there is no spread of fluorescence $\Delta F_r/\Delta F_0$ is expected to rapidly decay to zero away from the neuron membrane (assuming no signal from proximal dendrites). With the application of confocal gating to a targeted illumination microscope, we found that stronger confocal gating (smaller slit width) led to weaker $\Delta F_r/\Delta F_0$ across all measured distances up to 23.6 µm, with the drop being most significant just beyond the neuron membrane [Fig. S6(a), Table S4]. At larger distances, the differences in $\Delta F_r/\Delta F_0$ for different confocal slit sizes became smaller and less significant, particularly for smaller slit sizes of 4.5 and 11.3 µm ($p > 0.05$ for distance $\geq$ 14.2 µm, Table S4). Similarly, significant reductions in $\Delta F_r/\Delta F_0$ were also observed within 23.6 µm distances when targeted illumination was applied to a confocal microscope [Fig. S6(c)]. We therefore conclude that TICO microscopy is effective at reducing crosstalk from scattered fluorescence even in cases where the labeling is confined to a single layer (i.e. even in cases where there is no out-of-focus fluorescence), thus improving the fidelity of voltage imaging at high labeling density.

To quantify the added advantage of TICO microscopy in reducing out-of-focus background, we analyzed the correlations between the subthreshold membrane voltage (Vm) of neuron pairs throughout the imaging FOV. Neuronal populations tend to exhibit natural Vm correlations that are biological in origin; however, out-of-focus background can introduce additional apparent correlations that are erroneous. In principle, the ground truth associated with biological correlations could be obtained by pairwise electrophysiology, but such measurements are extremely difficult to perform, particularly in vivo, making it impossible to obtain sufficient statistics for generalizable results. We therefore adopted an indirect assessment of crosstalk, noting that biological Vm-Vm correlations should not depend on the imaging configuration (e.g. slit width, with/without targeted illumination, etc.), and that any observed configuration-induced changes in the measured Vm-Vm correlations must be the result of changes in crosstalk. We found that Vm-Vm correlations decreased significantly both when we decreased

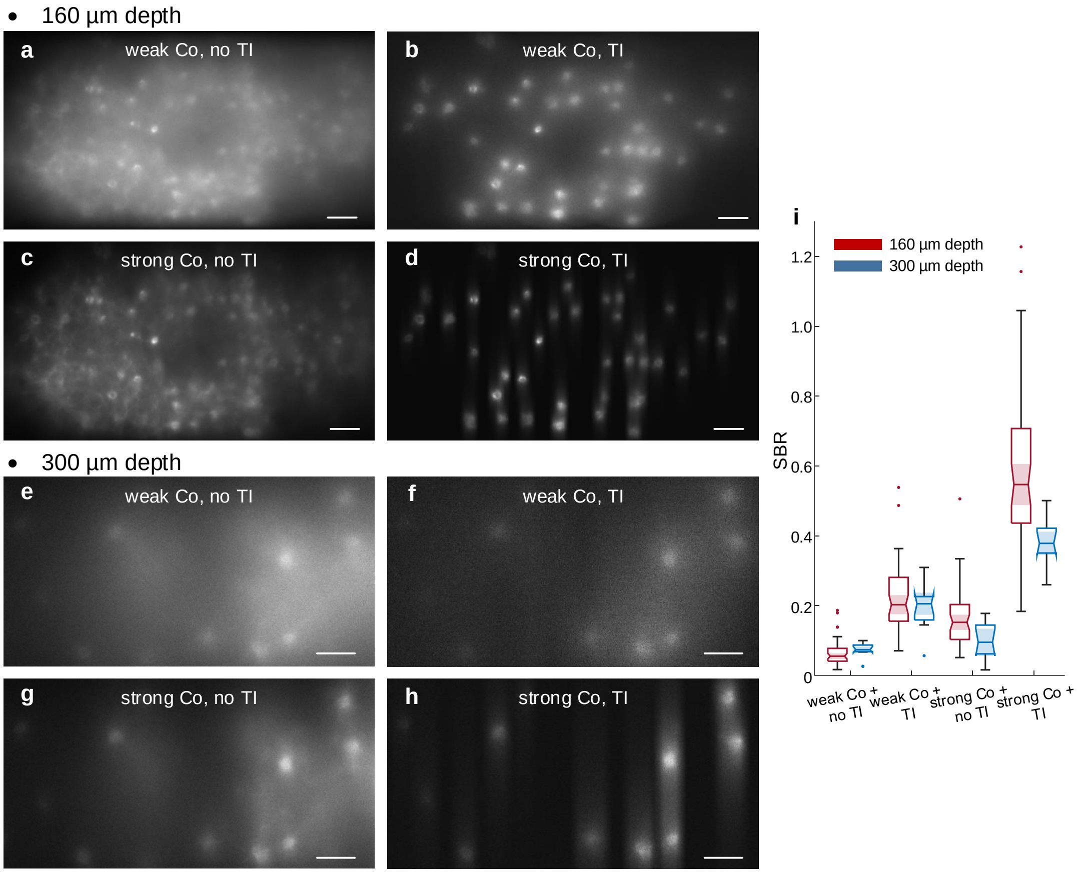


**Figure S5. Comparison of imaging contrast at different depths.**
(a-d) Voltron2 fluorescence imaged using TICO microscope under various combinations of weak/strong confocal detection (Co) and with/without targeted illumination (TI). Weak and strong confocal detection was achieved using a 156 μm and 14 μm confocal slit width respectively. Imaging depth at 160 μm. Scale bars are 50 μm.
(e-h) Same as (a-d) but imaged at depth 300 μm, with a strong confocal slit width set to 23 μm.
(i) Comparison of estimated SBR for different imaging depths and microscope configurations. For 160 μm depth, $n = 52$ cells; for 300 μm depth, $n = 11$ cells. Box plot the same as in Fig. 2(e) except that dots represent outliers. Scale bars in (a-h) are 50 μm.

confocal slit size and when we applied targeted illumination [Fig. S6(b,d), Table S4]. In addition, we observed similar Vm-Vm correlations beyond a pairwise distance of 200 μm when using smaller slits of 4.5 and 11.3 μm but not with the larger 22.5 and 156 μm slits sizes [Fig. S6(b), Table S4]. This indicates that moderate confocal gating (slit size $\sim$ 10 μm) is effective at rejecting far-out-of-focus fluorescence that contributes to spurious long-range correlations, but that stronger confocal gating is required if one wishes to remove near-out-of-focus fluorescence over short distances.

##### 2.3 TICO microscopy improves spike SNR and reduces photobleaching

Equally important for *in vivo* voltage imaging are SNR and photobleaching rates. These two parameters are interdependent and fundamentally different measures of microscope performance than SBR or $\Delta F/F$. The effects of targeted illumination and confocal gating are discussed below.

We begin by comparing spike SNR from the same neurons imaged under targeted illumination but with different confocal slit widths of 4.5, 11.3, 22.5, and 156 μm [6 FOVs from 2 mice]. As expected, increasing slit width allowed more signal to be captured thus leading to an increase in spike amplitude, but a net decrease in spike $\Delta F/F$ [Fig. 2(e)] owing to the more pronounced increase in background fluorescence. The overall balance of these trends determines the degree to which spikes can be distinguished from background noise. By evaluating the shot-noise-limited spike detection fidelity $d'$[6], we found this to be optimized near the intermediate slit width of 22.5 μm [Fig. 2(h)]. However, due to additional noise contributions inevitable to *in*

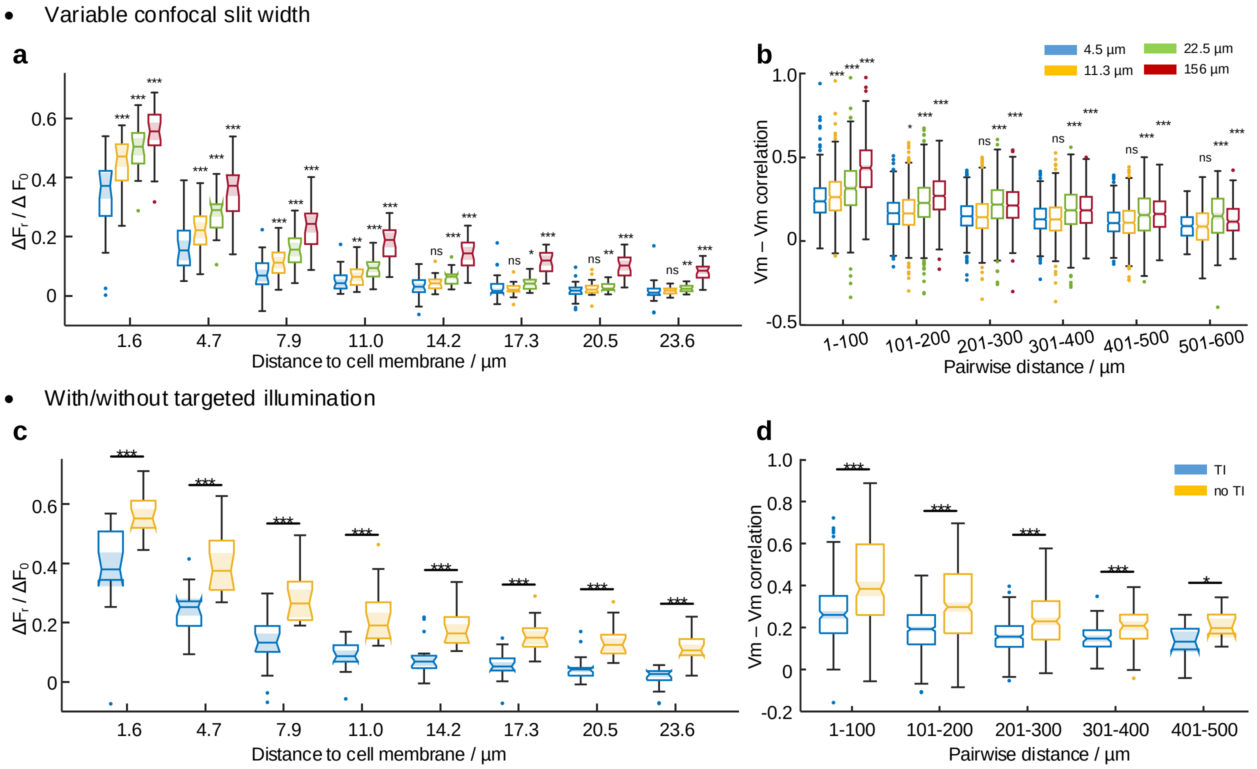


**Figure S6.** **Quantification of optical crosstalk under different microscope configurations.**
(a) Comparison of spike amplitude decay outside of cell membrane on a targeted illumination microscope with different confocal slit widths of 4.5, 11.3, 22.5, and 156 µm. n = 30 cells from 6 FOVs, 2 mice. Within each binned distance, "ns" not significant $p \geq 0.05$ , $*p < 0.05$, $**p < 0.01$, $***p < 0.001$, Wilcoxon signed-rank test compared to the control group of slit width 4.5 µm, see Table S4 for statistics. Box plot the same as in Fig. 2(e).
(b) Subthreshold Vm correlations between neuron pairs of varying separation. Boxplot same as (a) except that dots represent outliers. Within each binned distance, "ns" not significant, $***p < 0.001$, Wilcoxon signed-rank test compared to the control group of slit width 4.5 µm, see Table S4 for statistics.
(c,d) Same as (a,b) but comparing between with and without targeted illumination on a confocal microscope with 14 µm wide slit. n = 19 cells from 5 FOVs, 2 mice.

*vivo* imaging such as detection electronics, brain motion, hemodynamics, etc., we found smaller and less significant differences in the experimentally measured spike SNR for slit widths in the range 11.3 to 156 µm [Fig. 2(i), Table S4]. A significant reduction in spike SNR was observed only for the smallest 4.5 µm slit width, suggesting that shot noise was dominant only in this case, because of the small signal amplitude. These results are in qualitative agreement with the conclusions drawn from our theoretical modeling indicating that while achieving maximum SNR requires optimization of confocal gating strength, under *in vivo* imaging conditions this maximum is only weakly peaked and tolerant to a relatively wide range of slit widths (Fig. S2).

Interestingly, when applying targeted illumination to a confocal microscope (14 µm slit), we observed a small reduction in SNR from 5.66 to 5.13 [Fig. 2(j)] despite the lowered background shot noise, which is in contrast to the case for a widefield microscope where the application of targeted illumination led to higher SNR[7]. The main reason for the perceived SNR difference here was the reduction in excitation power delivered under targeted illumination. To explain in detail, neurons in scattering tissue can be excited directly by unscattered (ballistic) photons or indirectly by scattered photons. When the illumination is targeted to the neurons, the former remains unchanged whereas the latter can decrease significantly. This was observed experimentally from the reduction in photobleaching rate by 71.4% for neurons with targeted illumination compared to without [Fig. S7(a)], and also confirmed theoretically [Fig. S1(f)]. However, the SNR advantage of targeted illumination still remains. Because confocal gating preferentially detects signals produced by ballistic excitation, we observed a smaller reduction in baseline fluorescence of 60% [Fig. S7(b)]. Together with the 31.6% increase in spike contrast $\Delta F/F$ resulting from the stronger

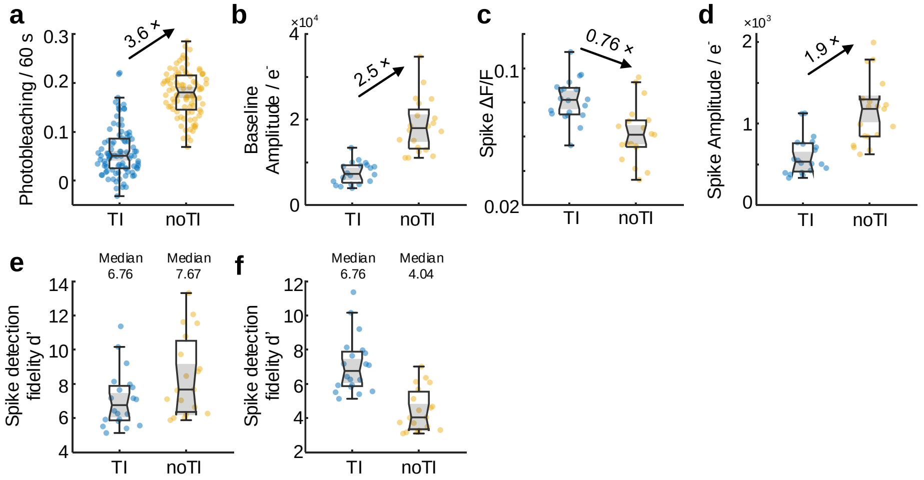


**Figure S7. Comparison of TICO microscope imaging performance with and without targeted illumination.**
(a-d) Comparison of photobleaching rate, baseline fluorescence amplitude, spike contrast $\Delta F/F$, and spike amplitude measured with and without targeted illumination (TI, noTI) with a 14 µm confocal slit. Same as main Fig. 2.
(e) Evaluation of theoretical shot-noise limited spike detection efficiency $d'$ according to the measured spike contrast $\Delta F/F$, and spike amplitude.
(f) Theoretical evaluation of spike detection efficiency $d'$ assuming equal photobleaching rate with and without targeted illumination. According to the measured reduction in photobleaching rate, the number of photoelectrons was reduced by 71.4% when without targeted illumination.

background rejection capacity [Fig. S7(c)], the overall decrease in spike amplitude under targeted illumination was only 47.4% [Fig. S7(d)], much less than the decrease in photobleaching rate. In fact, a theoretical evaluation of the configurations where targeted illumination is applied to a confocal versus a widefield microscope under the condition of equal photobleaching rate confirms that TICO microscopy provides higher spike detection fidelity [Fig. S7(e,f)], in accordance with the prediction from our simulation results (Fig. S3). We thus conclude that the experimentally observed reduction in SNR with targeted illumination was dominantly caused by the resulting reduction in scattered excitation power, though mitigated by the improved signal detection efficiency and background rejection provided by confocal gating. The reduced excitation power in turn led to a much lower photobleaching rate, providing a capacity for longer duration imaging.

#### 3 Evaluation of losses if the fluorescence were de-scanned through the DMD

A DMD chip consists of millions of micromirrors arranged in a 2D array, where each micromirror has two discrete tilt angles denoted by "On" and "Off". This periodic structure makes the DMD chip behave like a diffraction grating, where the incident light is diffracted into multiple diffraction orders with diffraction angle $\beta$ determined by the grating equation $p(\sin\alpha + \cos\beta) = m\lambda$, where $\alpha$ is the incident angle, $p$ is the grating pitch, $\lambda$ is the wavelength, and $m = ..., -1, 0, 1, ...$ is the diffraction order [Inset of Fig. S8(a)]. On the other hand, the angle of specular reflection, determined by the blaze angle $\theta_B$, is defined as $\beta' = -\alpha + 2\theta_B$. According to the blaze-angle condition, the highest diffraction efficiency can only be achieved when $\beta = \beta_B$ for a particular order. For fluorescence signals of large bandwidth, only certain wavelengths satisfy this blaze-angle condition. For other wavelengths, multiple diffraction orders must be collected in order to maximize transmission efficiency.

We consider a detection path of a fluorescence microscope where the fluorescent sample is imaged onto the DMD array surface and further re-imaged onto the camera. The detection aperture after the DMD [aperture A2 in Fig. S8(a)] determines the fluorescence collection efficiency, which is determined here by the mirror size of the galvanometer. Clearly, a larger-sized mirror would increase collection efficiency, but would also in turn introduce more inertia resulting in lower scan speed/angle. To ensure maximum frame rate and FOV, the mirror size should be matched to the back aperture size of the objective [aperture A1 in Fig. S8(a)]. As a result, except for certain diffraction orders at discrete wavelengths, the diffracted fluorescence suffers loss due to clipping by A2.

To further illustrate the effects of de-scanning through a DMD, we simulate the diffraction caused by the DMD and calculate the system transmission efficiency if the fluorescence signal were de-scanned through the DMD [Fig. S8(a)]. The DMD used in

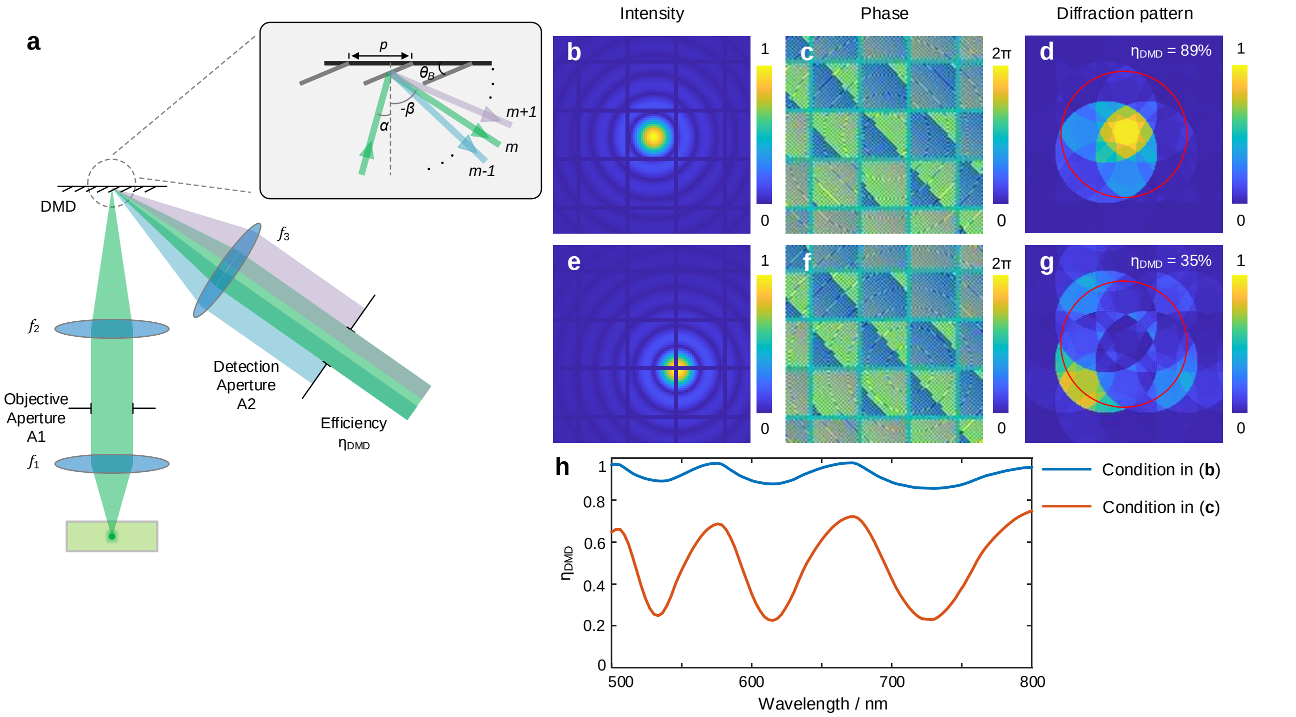


**Figure S8.** **Modeling of diffraction effect for fluorescence descanning through a DMD.**
(a) Schematic illustration of fluorescence descanning through a DMD.
(b, c) Intensity and phase profile when a fluorescent point source is imaged onto the DMD, the wavelength is assumed to be 600 nm.
(d) Corresponding far-field diffraction pattern at the detection aperture (red circle). The transmission efficiency through the detection aperture is 89%.
(e-g) Same as (b-d) but the fluorescent source is imaged to a different location of the DMD as shown in (e). The transmission efficiency now becomes 35%.
(h) Transmission efficiency $\eta_{DMD}$ as a function of wavelength for the two fluorescent source locations shown in (b,e).

our system has a mirror pitch $p = 13.68$ µm and tilt angle $\theta_B = \pm 12$ °. Based on our experimental setup, the focal length of each lens is $f_1 = 12.5$ mm, $f_2 = 180$ mm, $f_3 = 37.5$ mm, with the diameter of the objective aperture $A1 = 20$ mm, detection aperture $A2 = 5$ mm, and the NA of lens $f_1$ is 0.8. We use Fresnel propagation to compute the electric field distribution of a point fluorescent object when imaged onto the DMD, which is multiplied by the phase profile introduced by the DMD with all pixels assumed to be in "On" state, and then Fourier transformed by another lens onto the detection aperture A2. For more accurate results, we additionally include the finite DMD fill factor (92%), which leads to a dependence of the diffraction patterns on the fluorescent source location [Fig. S8(b-g)]. The ratio of the intensity within the detection aperture A2 to the total intensity at the A2 plane is used to determine the transmission efficiency (note that this does not include the finite fill factor of the DMD).

We calculate the DMD transmission efficiency over two spectral bands: 573 - 616 nm for Voltron imaging and 657 - 751 nm for somArchon imaging. The transmission efficiency is averaged across the wavelength bands (assuming flat spectra), and all possible source locations. Overall, the transmission efficiency would be 74% for Voltron imaging, and 63% for somArchon imaging. Accounting for additional losses due to the finite DMD fill factor (92%), DMD window transmission efficiency (96% double pass), and mirror reflectivity (89%)[8], the overall best transmission efficiencies if the fluorescence signals were descanned through the DMD would be 58% for Voltron, and 49% for somArchon.

#### 4 Derivation of the spike detection fidelity obtained with a scanning microscope

To calculate the theoretical shot-noise-limited spike detection fidelity $d'$, we follow the procedure outlined in Ref.[6]. In detail, we assume a fluorescence signal model given by

$$F(t \geq 0) = F_0 + F_{AP} \cdot e^{-(t-t_0)/\tau} \tag{S17}$$

where $F_0$ is the baseline fluorescence, $F_{AP}$ is the spike amplitude, $\tau$ is the decay constant of the fluorescent indicator, $t_0 \in [-1/v, 0]$ is the onset time of the spike event, and $v$ is the sampling rate of the imaging system. In our case, as in the case for most voltage imaging microscopes, because the sampling interval is comparable to the fluorescence decay time, we assume a single time point at $t = 0$ is used to detect spike events. For a scanning microscope, the excitation intensity is inversely proportional to the integration time $1/\varphi v$. With an infinitely small integration time $\varphi \to 0$ (a scanning microscope), the detected fluorescence signal and background at time $t = 0$ can be written as, respectively:

$$S_0 = \lim_{\varphi \to 0} \frac{1}{\varphi v} \int_0^{\varphi} F(t)dt = \frac{F_0}{v} + \frac{F_{AP}}{v} \cdot e^{t_0/\tau} \tag{S18}$$

and

$$B_0 = \lim_{\varphi \to 0} \frac{1}{\varphi v} \int_0^{\varphi} F_0 dt = F_0/v \tag{S19}$$

Therefore, making use of Poisson statistics, the probabilities associated with obtaining a single measurement of photon number $N$ without a spike event $H^{(0)}$ and with a spike event $H^{(1)}$ can be written as:

$$p(N|H^{(0)}) = B_0^N e^{-B_0}/N! \tag{S20}$$

$$p(N|H^{(1)}) = S_0^N e^{-S_0}/N! \tag{S21}$$

leading to a log-likelihood ratio

$$L(N) = \log \frac{p(N|H^{(1)})}{p(N|H^{(0)})} = N \log \frac{S_0}{B_0} - S_0 + B_0 \tag{S22}$$

With the assumption that $\frac{F_{AP}}{F_0} \ll 1$, we can calculate the mean $\mu_L^{(1,0)}$ and variance $\sigma_L^{(1,0)}$ of $L(N)$ under the assumptions of a spike occurring or not:

$$\mu_L^{(0)} = \frac{F_0}{v} \log(1 + \frac{F_{AP}}{F_0} e^{t_0/\tau}) - \frac{F_{AP}}{v} \cdot e^{t_0/\tau} \approx -\frac{F_{AP}^2}{2F_0 v} e^{2t_0/\tau} \tag{S23}$$

$$\mu_L^{(1)} = \frac{1}{v}(F_0 + F_{AP} \cdot e^{t_0/\tau}) \log(1 + \frac{F_{AP}}{F_0} e^{t_0/\tau}) - \frac{F_{AP}}{v} \cdot e^{t_0/\tau} \approx \frac{F_{AP}^2}{2F_0 v} e^{2t_0/\tau} \tag{S24}$$

$$(\sigma_L^{(0)})^2 = \frac{F_0}{v} \log^2(1 + \frac{F_{AP}}{F_0} e^{t_0/\tau}) \approx \frac{F_{AP}^2}{F_0 v} e^{2t_0/\tau} \tag{S25}$$

$$(\sigma_L^{(1)})^2 = \frac{1}{v}(F_0 + F_{AP} \cdot e^{t_0/\tau}) \log^2(1 + \frac{F_{AP}}{F_0} e^{t_0/\tau}) \approx \frac{F_{AP}^2}{F_0 v} e^{2t_0/\tau} \tag{S26}$$

Following the same definition of spike detection fidelity index[6] $d' = (\mu_L^{(1)} - \mu_L^{(0)})/\sigma_L^{(0)}$, we have $d'(t_0) = \sqrt{\frac{F_{AP}^2}{F_0 v} e^{2t_0/\tau}}$. If the spike onset time $t_0$ is distributed uniformly over $[-1/v, 0]$, the averaged $d'$ is found to be:

$$\bar{d'} = v \int_{-1/v}^{0} d'(t_0) dt_0 = \tau v (1 - e^{-1/\tau v}) \frac{F_{AP}}{\sqrt{F_0 v}} \tag{S27}$$

$F_{AP}$ can be obtained from the experimentally measured average $\Delta F/F$ according to:

$$\Delta F/F = \frac{\int_{-1/v}^{0} \frac{F_{AP}}{F_0} e^{t_0/\tau} dt}{1/v} = \tau v (1 - e^{-1/\tau v}) \frac{F_{AP}}{F_0} \tag{S28}$$

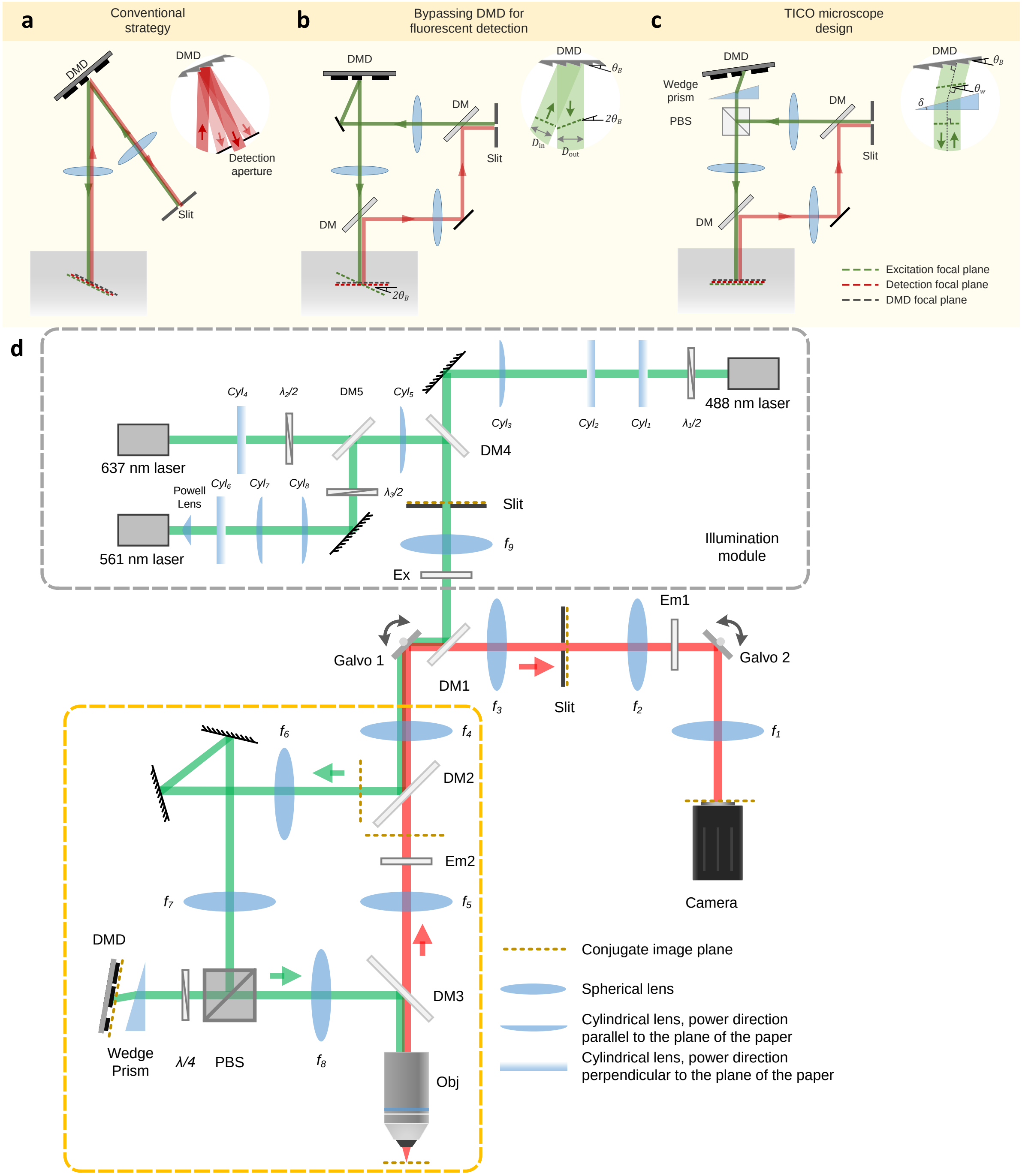


**Figure S9. Principle and schematic of TICO microscope.**
(a-c) Design principle of TICO microscope: (a) conventional strategy for incorporation a DMD into a confocal microscope; (b) bypassing DMD in the detection path to avoid fluorescence loss; (c) inserting a wedge prism in front of the DMD corrects for both image plane tilt and 1D magnification change caused by the DMD, restoring confocality between excitation and detection beams.
(d) Detailed schematic of TICO microscope. DM, dichromatic mirror. Em, emission filter. Ex, excitation filter. PBS, polarizing beam splitter. $\lambda/2$, half-wave plate. $\lambda/4$, quarter-wave plate. Galvo, galvanometric scanner. DMD, digital micromirror device. Obj, objective.

.

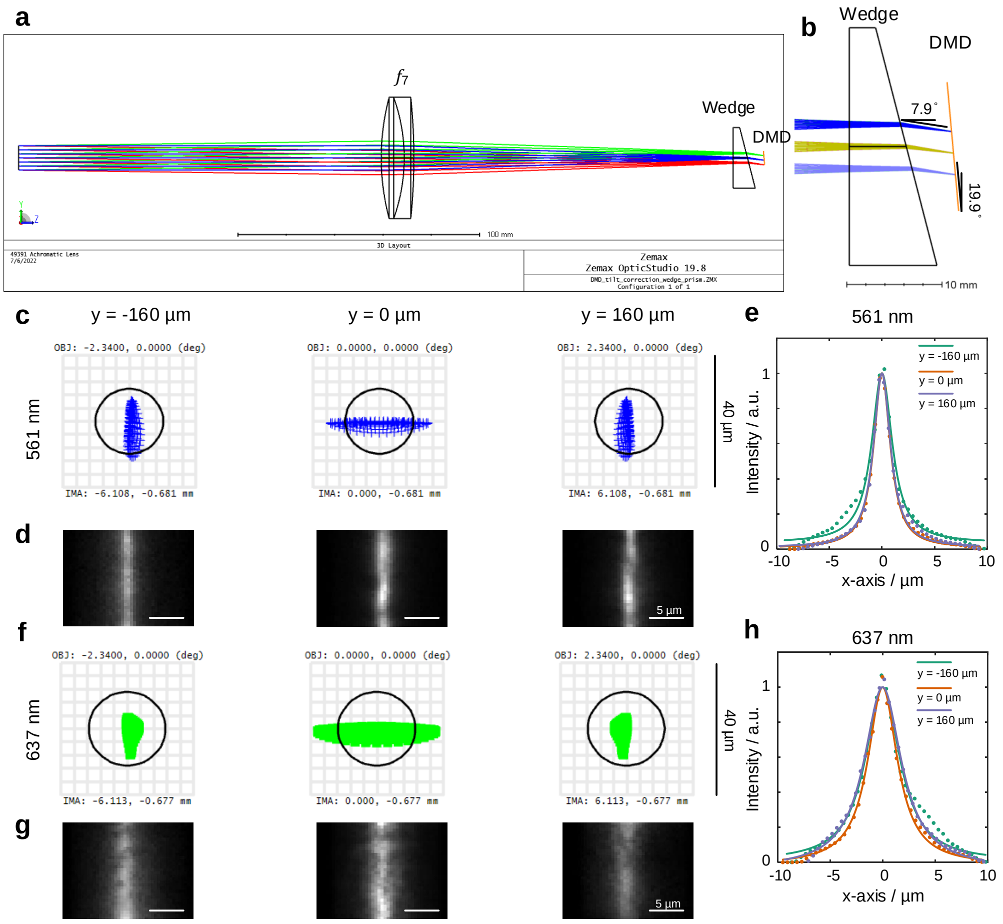


**Figure S10.** **Zemax simulation of focusing on a tilted image plane through a wedge prism.**
(a) Zemax cross-sectional view of the system. The input aperture was set to 10 mm, corresponding to 0.4 NA in the object space. Focusing lens $f_7 = 150$ mm, Thorlabs AC508-150-A. Wedge prism made of N-BK7 glass with apex angle 14°51', Edmund Optics 49-443. Image plane (DMD plane) tilted at 19.9°.
(b) Zoomed-in view at the wedge prism showing matched tilt angle at the DMD surface (19.9°- 7.9 °= 12 °corresponding to the micromirror tilt angle).
(c) Zemax spot diagram for 561 nm wavelength at different vertical positions corresponding to object space locations $y = -160, 0, 160$ µm.
(d) Image of the 561 nm laser line focus of the TICO microscope captured from the camera at object space locations $y = -160, 0, 160$ µm.
(e) Cross-sectional intensity profiles of the line foci shown in (d). Dots, measurement points; continuous lines, Lorentzian fit.
(f-h) Same as (c-e) but for 637 nm wavelength.

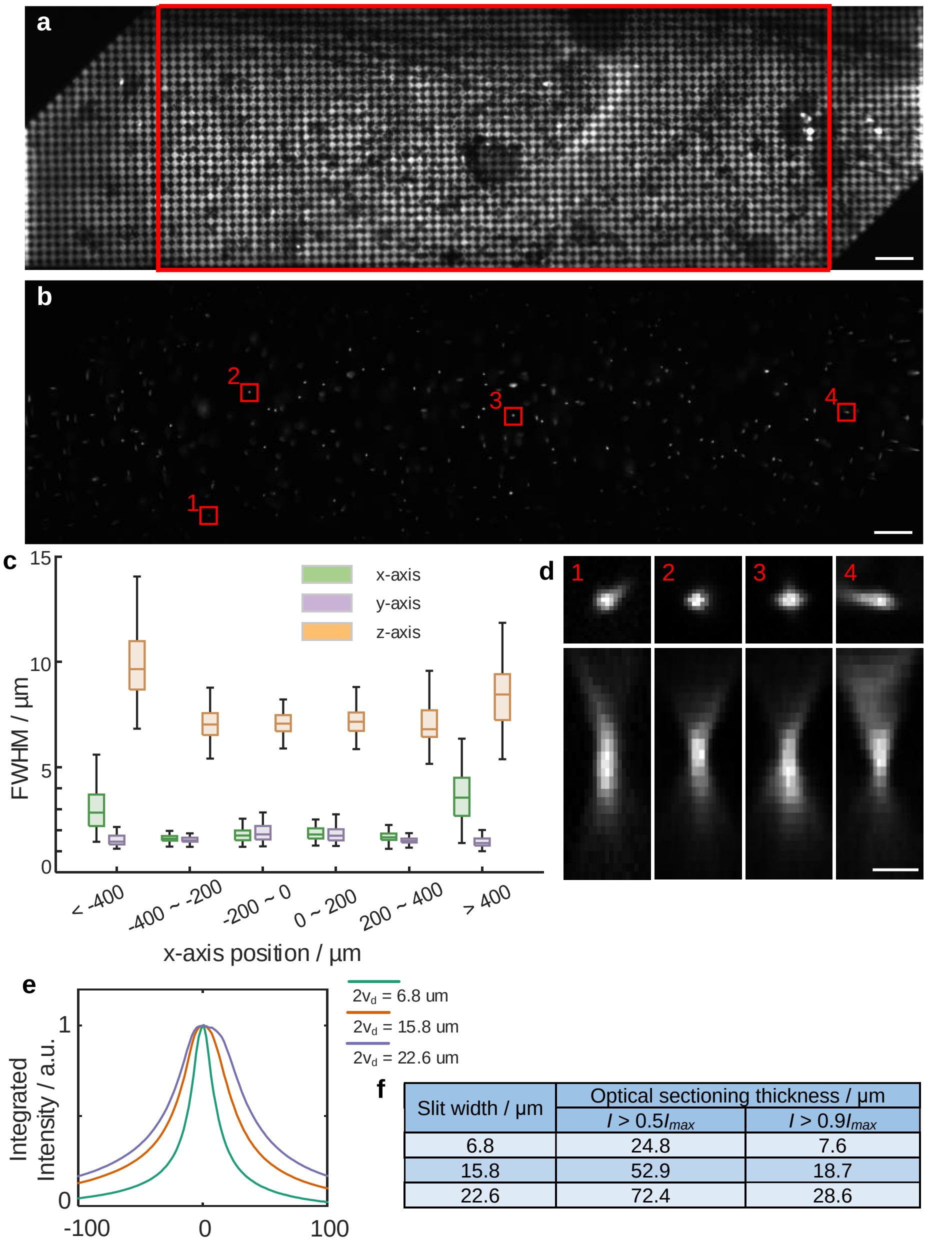


| Slit width / µm | Optical sectioning thickness / µm | |
|---|---|---|
| | $I > 0.5I_{max}$ | $I > 0.9I_{max}$ |
| 6.8 | 24.8 | 7.6 |
| 15.8 | 52.9 | 18.7 |
| 22.6 | 72.4 | 28.6 |

**Figure S11. Optical performance characterization of TICO microscope.**
(a) Fluorescence image of a single layer of 1 µm fluorescent beads acquired by projecting a 7.9 µm checkerboard pattern on the DMD. Note that over the full FOV of 1.16 × 0.325 mm the top left and bottom right corner are clipped due to the smaller DMD chip size. The FOV without clipping is 880 × 325 µm, indicated by the red rectangle. Image shows sufficient resolution for soma targeting across the entire FOV. Scale bar, 50 µm.
(b) Confocal image of 100 nm fluorescent beads over the FOV. Slit size was set to 14 µm.
(c) FWHM values of PSFs across different lateral positions across the FOV.
(d) Example PSFs from the red rectangular regions shown in (b).
(e) Optical sectioning profiles measured with different slit widths $2v_d$. Data obtained by axially translating a single layer of 1 µm fluorescent beads and measuring the integrated intensity as a function of defocus without targeted illumination. a.u., arbitrary unit.
(f) Thickness of optical sections measured at a threshold of 50% or 90% of the maximum intensity.

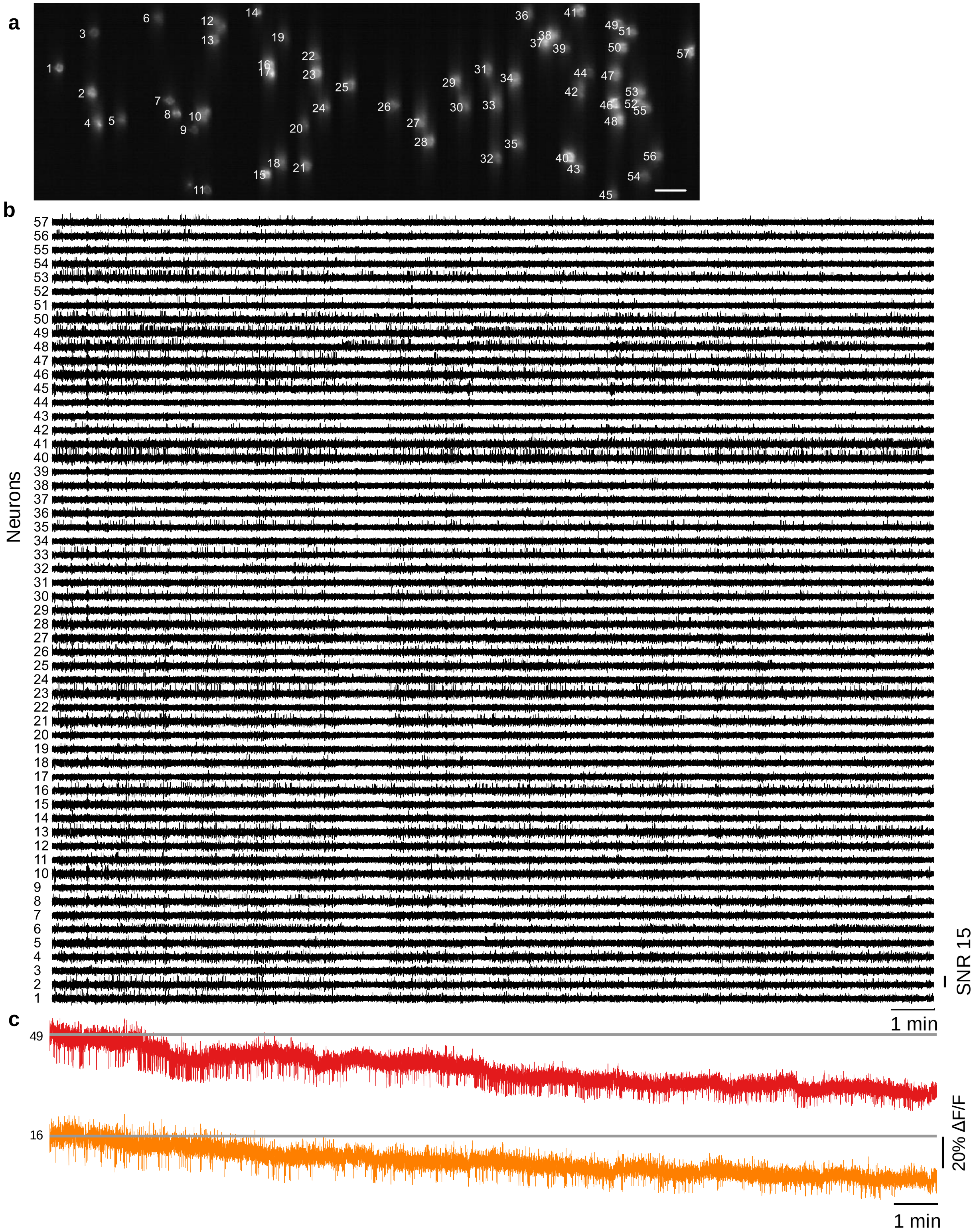


**Figure S12.** **Large-scale imaging of Voltron2 fluorescence from 57 cells in vivo.**
(a) Averaged Voltron2 fluorescence image from TICO microscope with 57 cells targeted. Scale bar, 50 µm.
(b) Complete 20 min recording of Voltron2 fluorescence from 57 cells.
(c) Raw fluorescence traces from 2 selected cells.

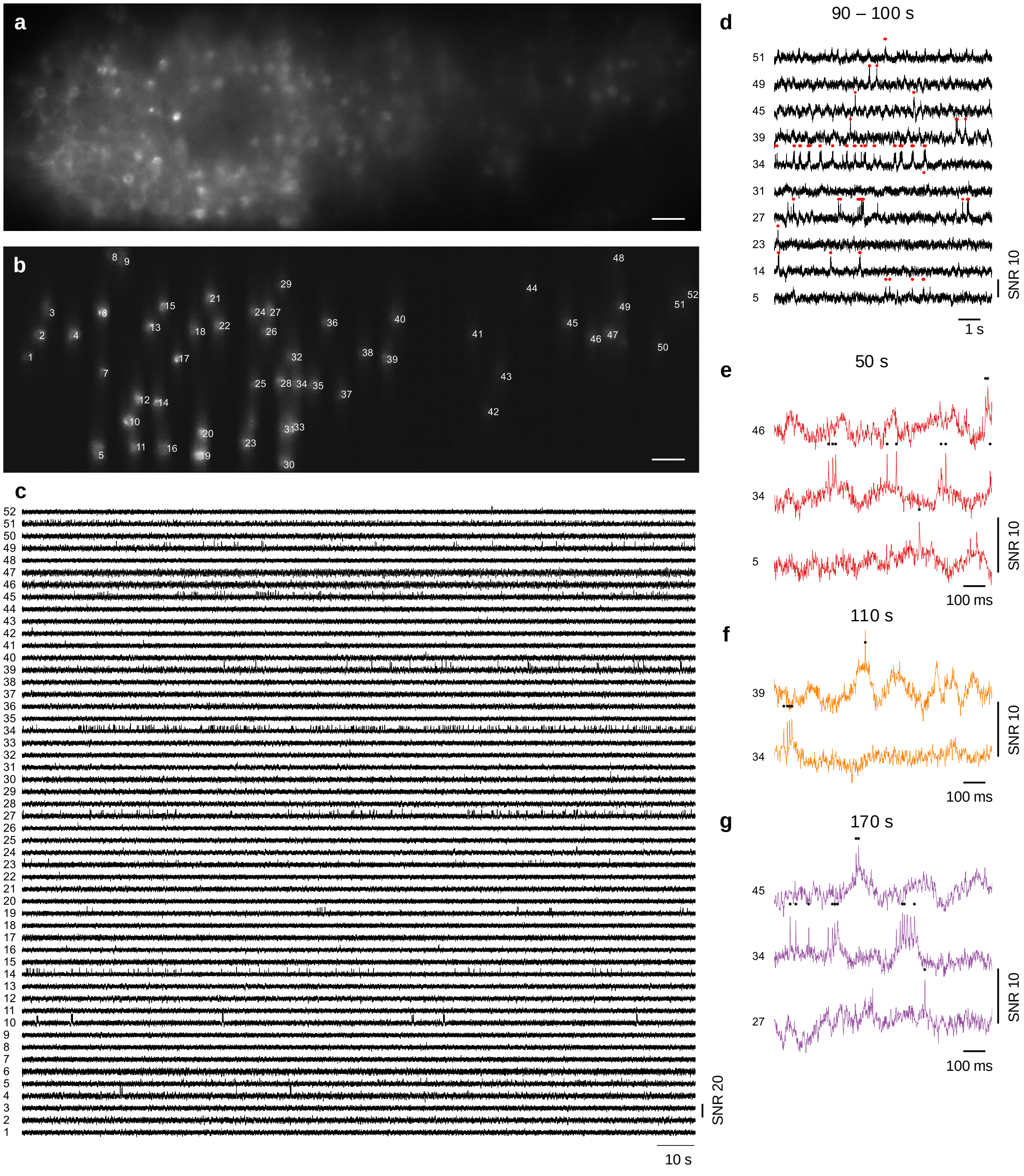


**Figure S13.** **Large-scale imaging of Voltron2 fluorescence from 52 neurons in vivo.**
(a) Confocal image of Voltron2 fluorescence over the imaging FOV. Scale bar, 50 μm.
(b) Average Voltron2 fluorescence image with 52 neurons targeted within the FOV. Scale bar, 50 μm.
(c) Voltron2 fluorescence traces from all 52 neurons over a 3 min recording.
(d) Zoomed-in fluorescence traces from 10 active cells during the recording period between 90 - 100 s. Dots, spike locations.
(f-g) Further zoomed-in fluorescence traces from active cells at recording times 50 s, 110 s and 170 s.

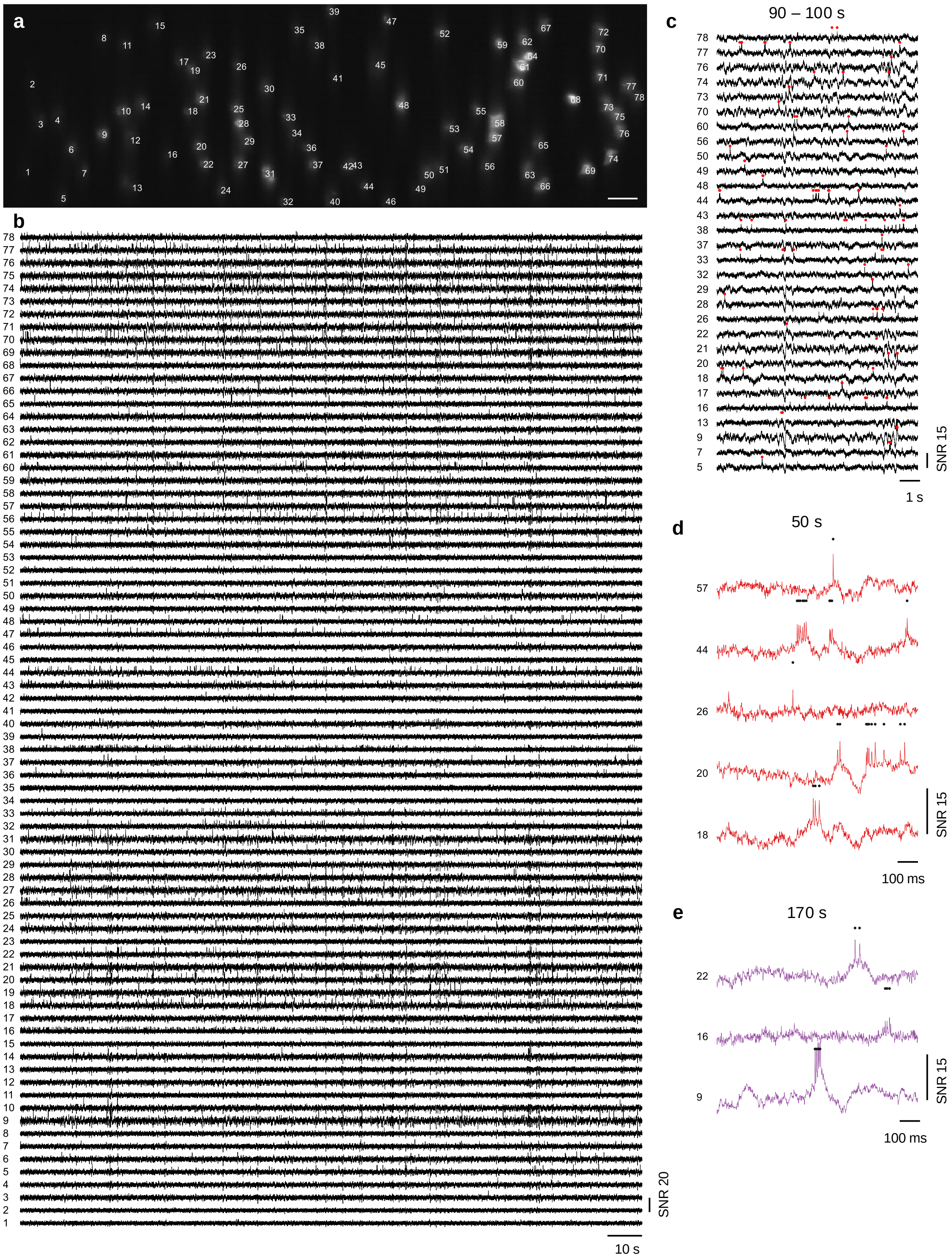


**Figure S14.** **Large-scale imaging of Voltron2 fluorescence from 78 neurons in vivo.**

**Figure S14.** (a) Average Voltron2 fluorescence image. Scale bar, 50 µm.
(b) Voltron2 fluorescence traces from all 78 neurons over a 3 min recording.
(c) Zoomed-in fluorescence traces from 30 active neurons during the recording period between 90 - 100 s. Dots, spike locations.
(d,e) Further zoomed-in fluorescence traces from active cells at recording times 50 s and 170 s.

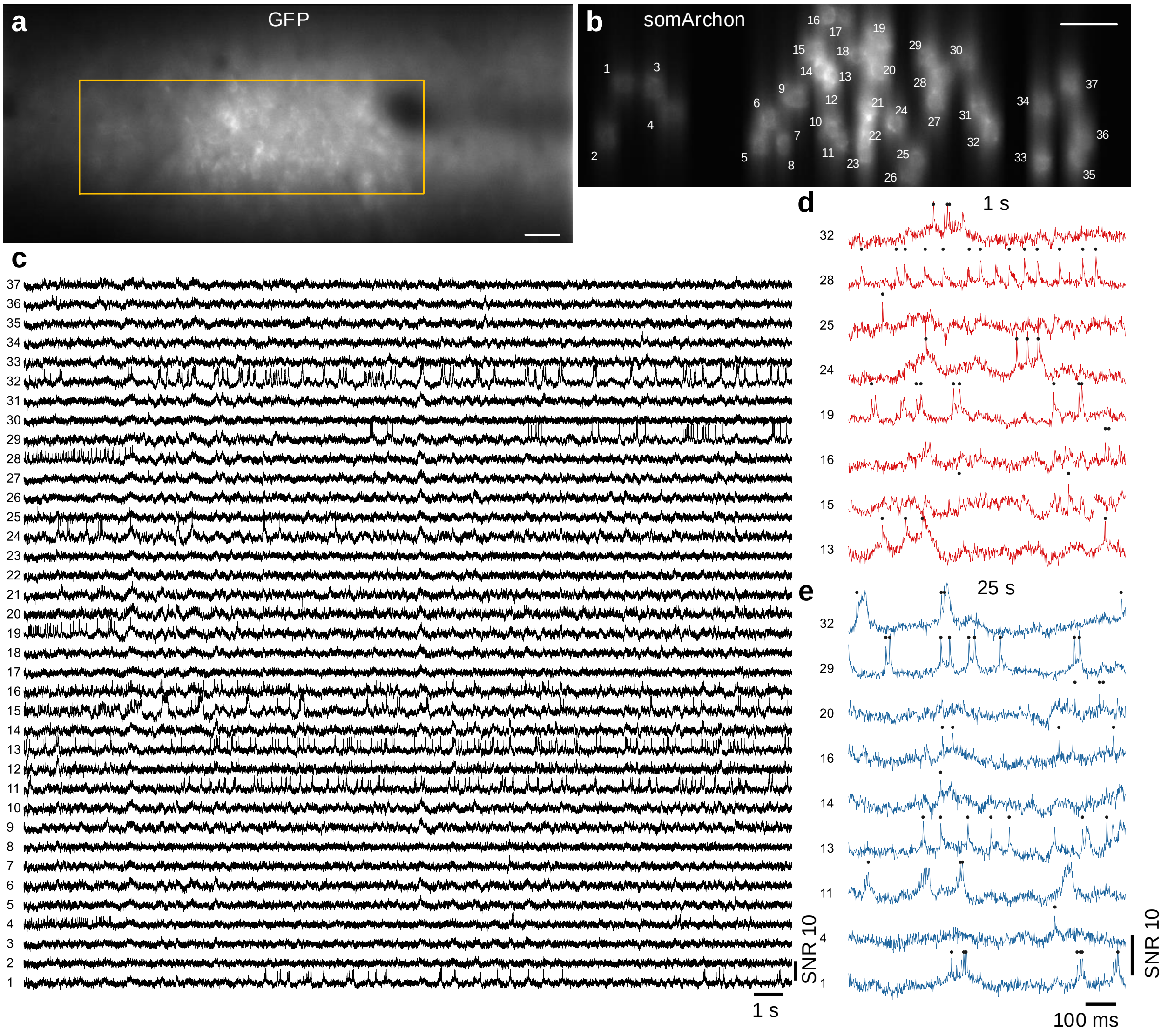


**Figure S15. Large-scale imaging of somArchon fluorescence from 37 cells near the visual cortex.**
(a) Confocal image of GFP fluorescence. Yellow square indicates actual somArchon imaging FOV shown in (b). Scale bar, 50 µm.
(b) SomArchon fluorescence image with 37 cells targeted. Scale bar, 50 µm.
(c) SomArchon fluorescence traces of 37 cells over a continuous 30 s recording. Recording speed 775 Hz, imaging depth 100 µm.
(d,e) Zoomed-in fluorescence traces of active neurons during 1 s and 25 s of the recording.

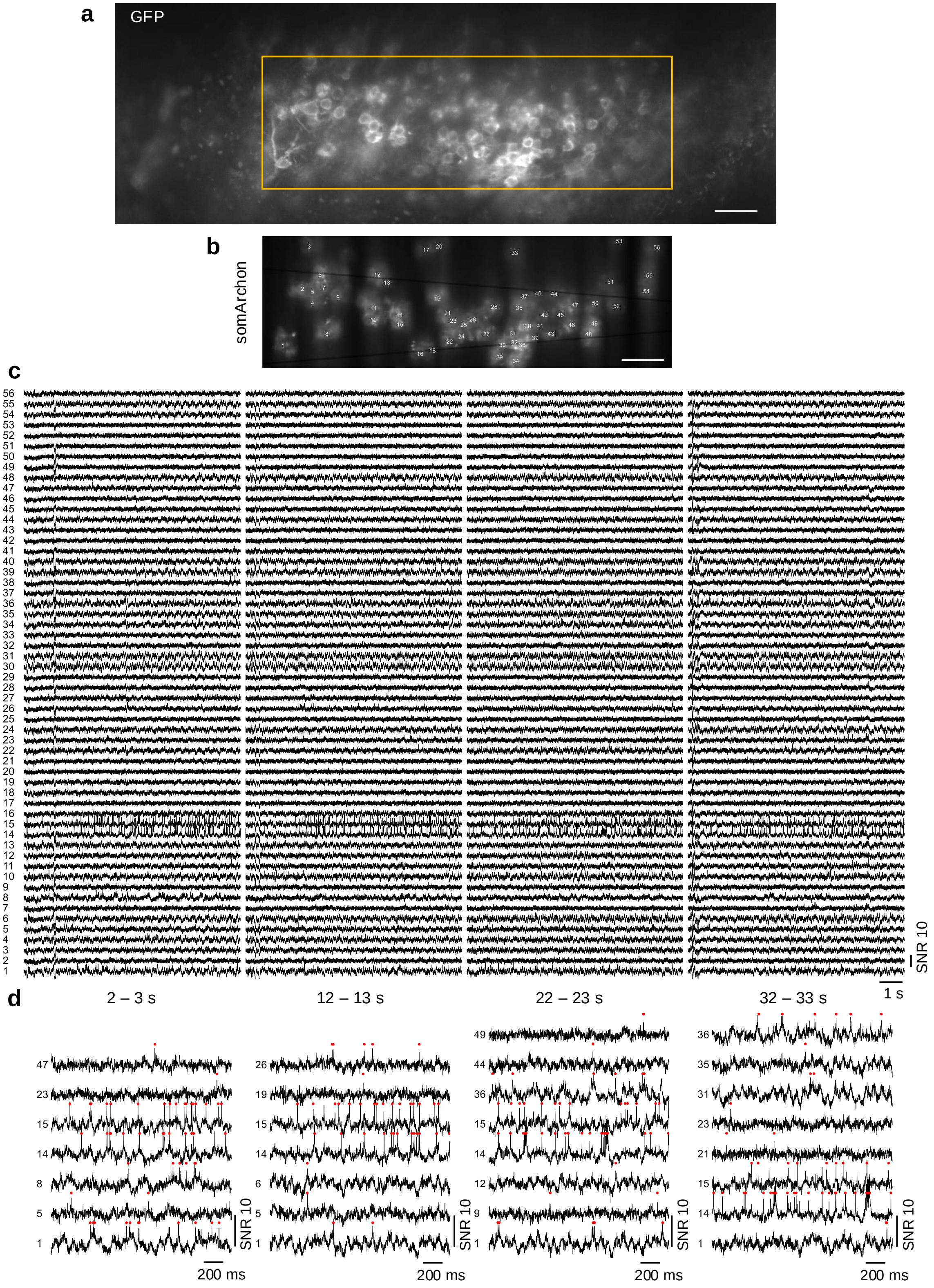


**Figure S16.** **Large-scale imaging of somArchon fluorescence from 56 cells in the hippocampus.**

**Figure S16.** (a) Confocal image of GFP fluorescence. Yellow square indicates actual somArchon imaging FOV shown in (b). Scale bar, 50 µm.
(b) SomArchon fluorescence image with 56 cells targeted within the FOV. Scale bar, 50 µm.
(c) SomArchon fluorescence traces of 56 cells over 4 separate 10 s recordings. Recording speed 800 Hz, imaging depth 80 µm.
(d) Zoomed-in fluorescence traces of active neurons during 2 s clips.

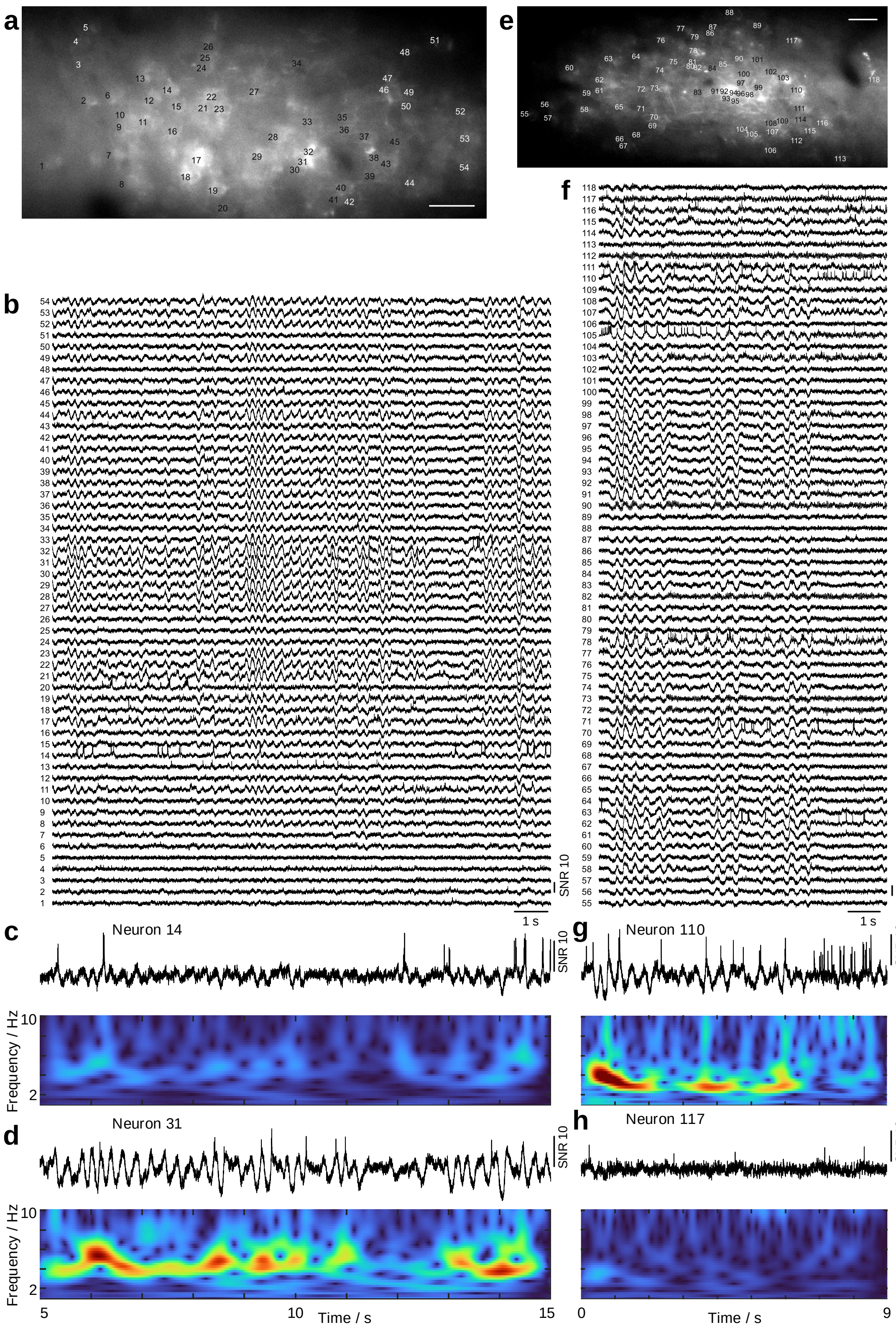


**Figure S17.** **Observation of highly synchronized 3 - 5 Hz membrane oscillations in L1 interneurons.**

**Figure S17.** (a) Confocal image of GFP fluorescence over the imaging FOV.
(b) SomArchon fluorescence traces for all the neurons labeled in (a).
(c,d) Zoomed-in fluorescence traces (top panel) of two selective neurons and their corresponding power spectra (bottom panel).
(e-h) Same as (a-d) but for a different FOV.

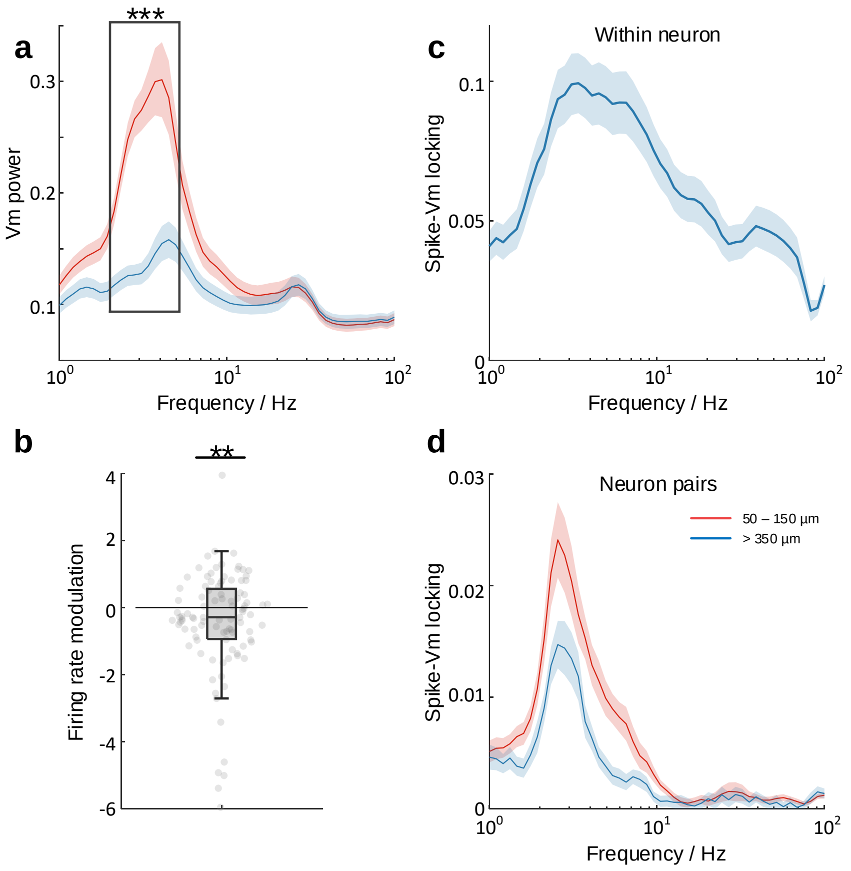


**Figure S18. Analysis of voltage traces in Fig. S17.**
(a) Frequency-resolved Vm power averaged from time periods with high delta population Vm power (> 2 standard deviation, S.D.; red trace) and low Vm power (< 2 S.D.; blue trace). Within 2 - 5 Hz frequency range (black box), most neurons showed significant Vm delta power modulation (paired student t-test, $***p = 3.27e^{-12}$, n = 99 neurons with average spike rate $\geq$ 1 Hz, 8 FOVs from 1 mouse). Shaded area, $\pm$1 S.D.
(b) Firing rate modulation of neurons from periods of high Vm delta power relative to periods of of low Vm delta power. Paired student t-test, $**p = 0.009$, n = 99 neurons with average spike rate $\geq$ 1 Hz, 8 FOVs from 1 mouse.
(c) Frequency-resolved spike-Vm phase locking for all neurons. Shaded area, $\pm$1 S.D.
(d) Frequency-resolved pike-Vm phase locking between neuron pairs. Red trace, neuron pairs with separation distances between 50 - 150 µm; blue trace, neuron pairs with separation distances > 350 µm. Shaded area, $\pm$1 S.D.

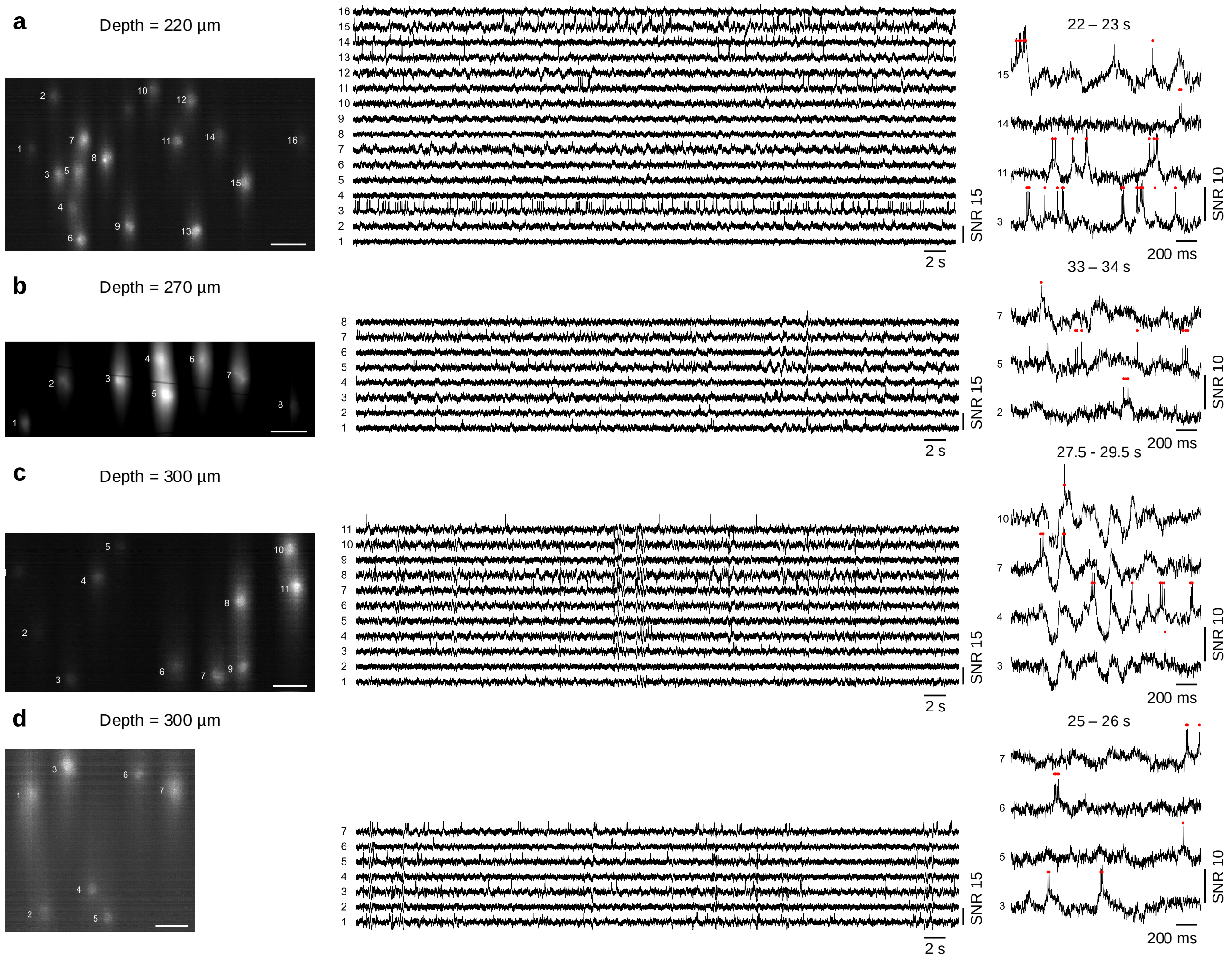


**Figure S19. Additional datasets for *in vivo* imaging at depths greater than 200 µm.** Imaging depths from (a-d) are 220, 270, 300, and 300 µm. Left column, averaged Voltron2 fluorescence image. Scale bars are 50 µm. Middle column, Voltron2 fluorescence traces from corresponding labeled neurons over 60 s recordings. Right column, zoomed-in fluorescence traces of active neurons during 2 s clips.

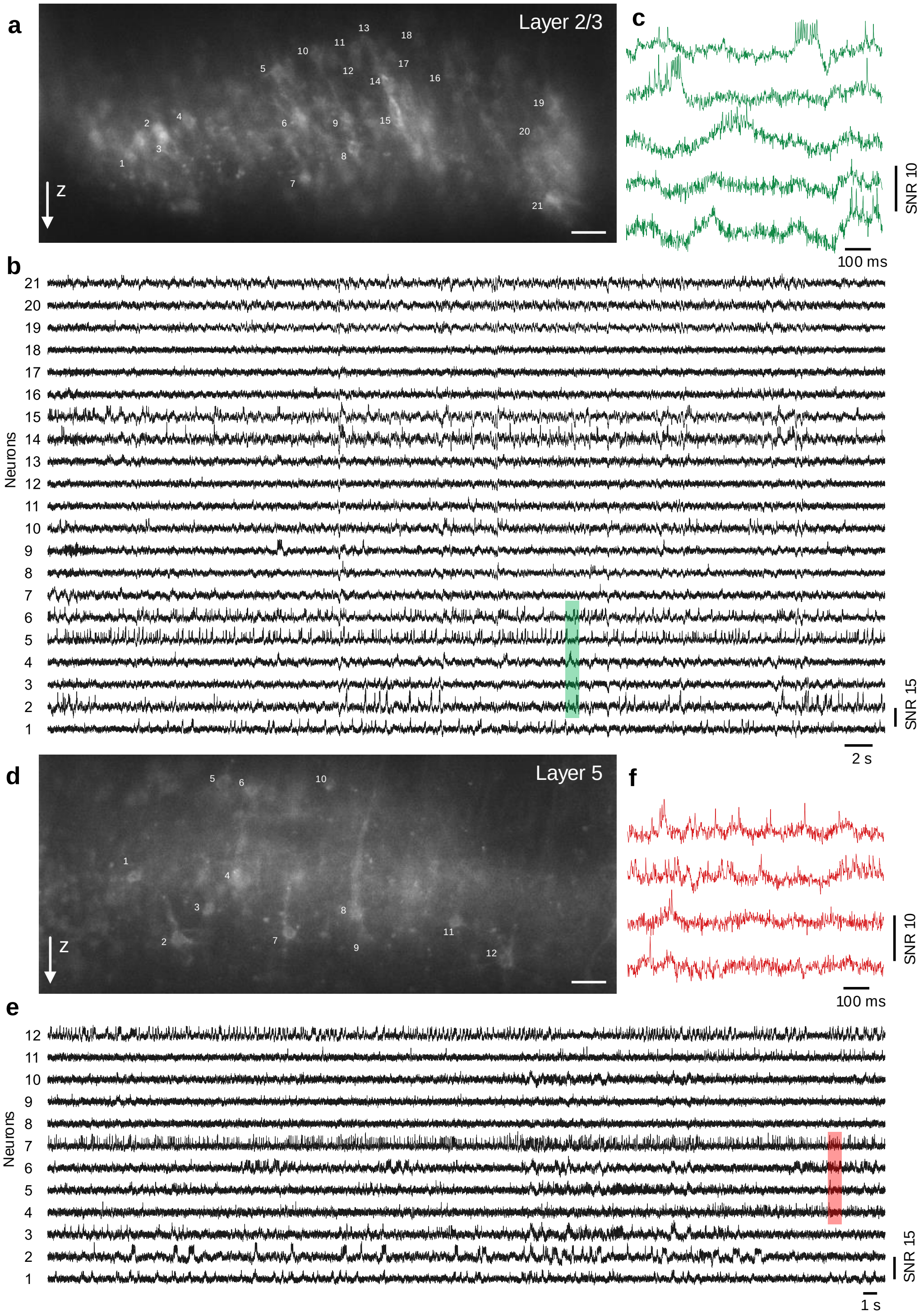


**Figure S20. Deep cortical voltage imaging via an implanted microprism.**
(a) Confocal image of Voltron2 fluorescence imaged via an implanted microprism over cortical layer 2/3. Scale bar, 50 μm.
(b) Voltron2 fluorescence traces of the 21 targeted neurons over a continuous 60 s recording. Recording speed 800 Hz.
(c) Zoomed-in fluorescence traces over the shaded area in (b).
(d-e) Same as (a-c) but imaged over cortical layer 5.

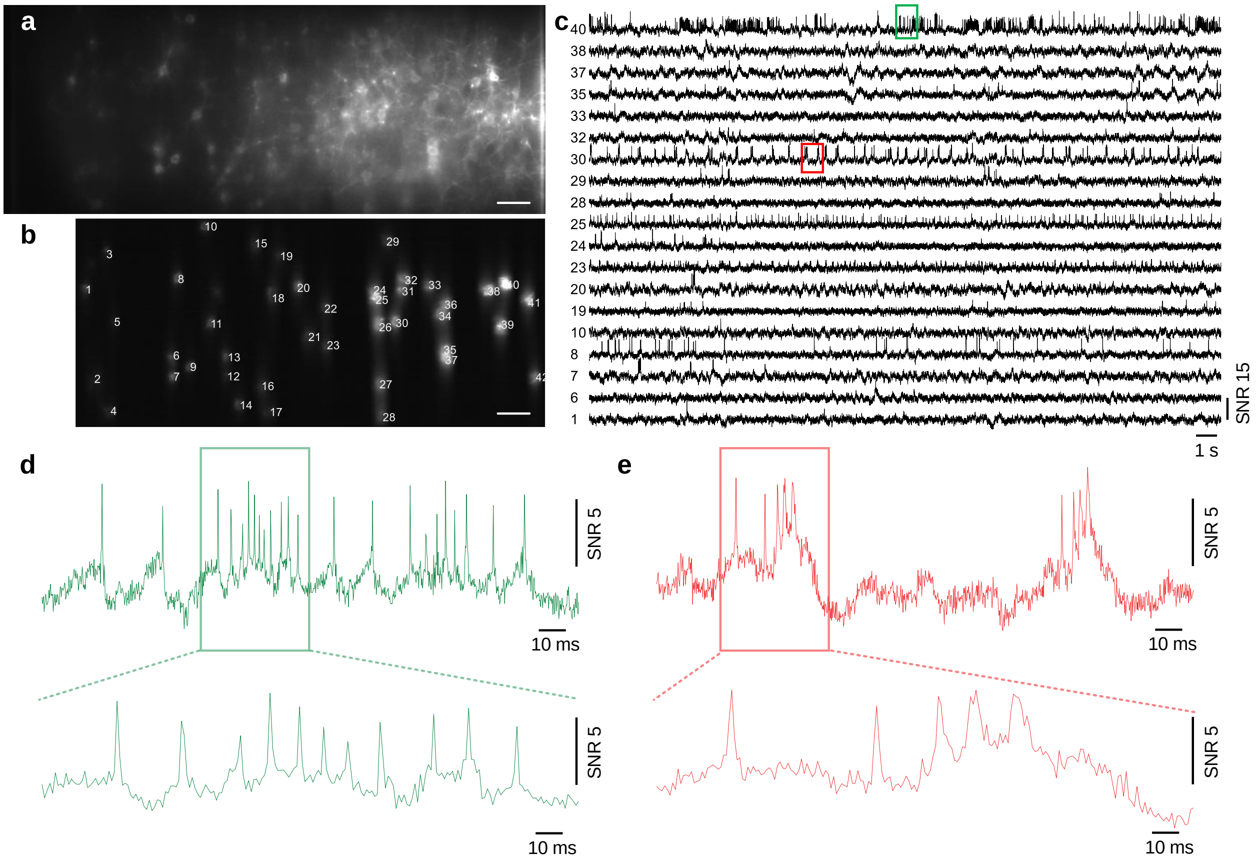


**Figure S21.** **High-speed voltage imaging at 1 kHz frame rate.**
(a) Confocal image of Voltron2 fluorescence over a FOV of 880 × 325 µm. Scale bar, 50 µm.
(b) Averaged Voltron2 fluorescence image with 42 targeted neurons.
(c) Fluorescence traces of spiking neurons over a 30 s recording.
(d,e) Zoomed-in fluorescence traces over the rectangular labeled regions in (c).

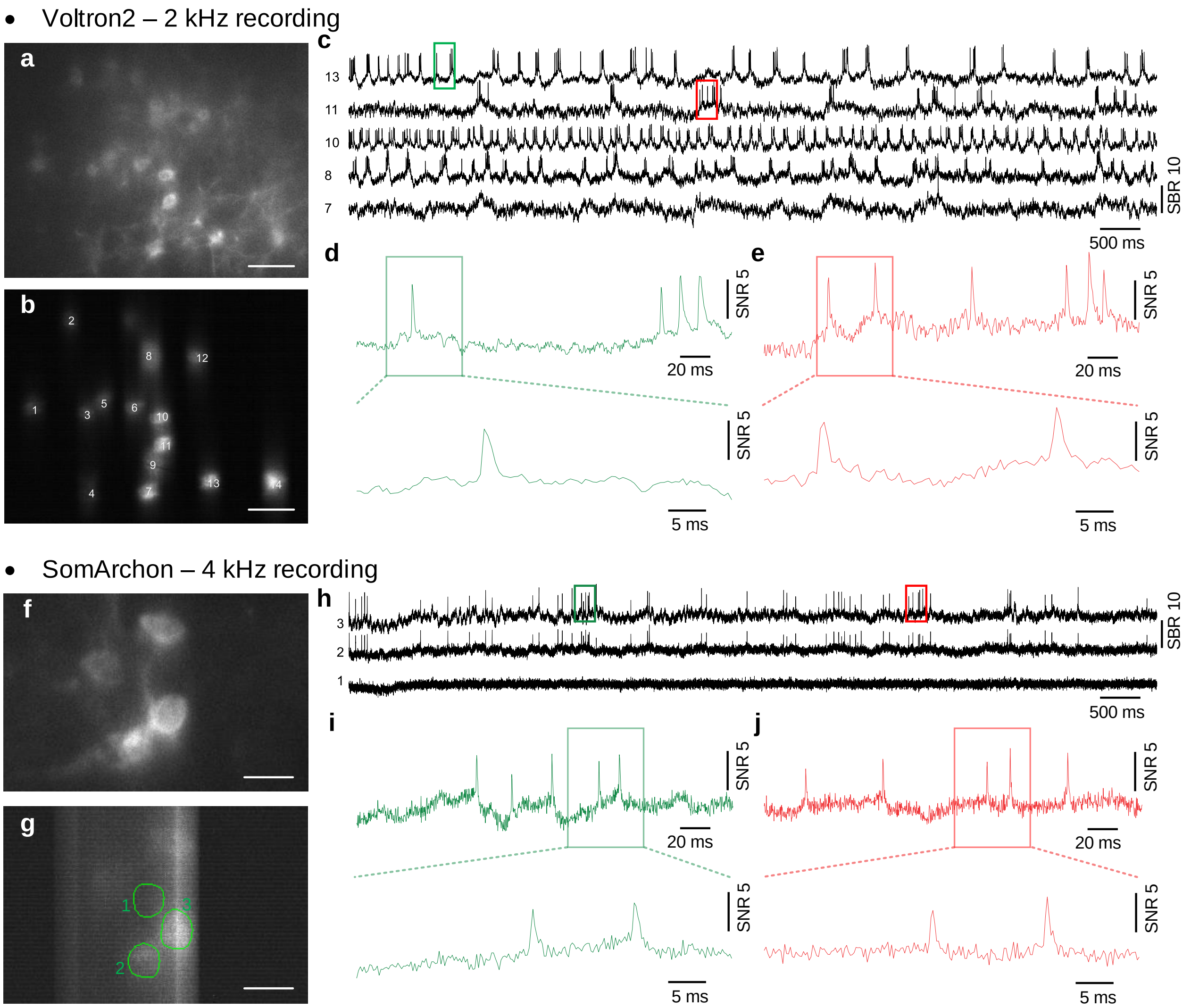


**Figure S22. High-speed voltage imaging at 2 kHz and 4 kHz frame rates.**
(a) Confocal image of Voltron2 fluorescence. Scale bar, 50 µm.
(b) Averaged Voltron2 fluorescence image with 14 neurons targeted within the FOV. Scale bar, 50 µm.
(c) Voltron fluorescence traces of 5 active neurons over a 10 s recording. Recording speed 2 kHz.
(d,e) Zoomed-in fluorescence traces over the rectangular labeled regions in (c).
(f) Confocal image of GFP fluorescence. Scale bar, 20 µm.
(g) Averaged somArchon fluorescence image with 4 neurons targeted within the FOV. Scale bar, 20 µm.
(h) SomArchon fluorescence traces of 2 active neurons over a 10 s recording. Recording speed 4 kHz. Note that the identical spiking activities observed in neuron 2 and 3 were caused by physiological reasons and not by optical crosstalk, since a nearby ROI 1 next to neuron 3 did not exhibit such activities.
(i,j) Zoomed-in fluorescence traces over the rectangular labeled regions in (h).

**Table S2.** Full list of components used in TICO microscope.

| Component | Description | Manufacturer | Part number |
|---|---|---|---|
| $f_1, f_4$ | 37.5 mm focal length Plossl eyepiece | Thorlabs | $2\times$ AC254-075-A |
| $f_2, f_3, f_9$ | 100 mm focal length achromatic doublet | Thorlabs | AC254-100-A |
| $f_5, f_6$ | 180 mm focal length achromatic doublet | Thorlabs | AC508-180-AB |
| $f_7, f_8$ | 150 mm focal length achromatic doublet | Thorlabs | AC508-150-A |
| $Cyl_1, Cyl_7$ | -50 mm focal length cylindrical lens | Thorlabs | LK1336RM-A |
| $Cyl_2, Cyl_3, Cyl_8$ | 150 mm focal length cylindrical lens | Thorlabs | LJ1629RM-A |
| $Cyl_4, Cyl_5$ | 100 mm focal length cylindrical lens | Thorlabs | LJ1567RM-A |
| $Cyl_6$ | 75 mm focal length cylindrical lens | Thorlabs | LJ1703RM-A |
| Wedge prism | N-BK7 14°51' apex angle wedge prism | Edmund Optics | 49-443 |
| Powell Lens | Powell lens | Laserline Optics Canada | LOCP-8.9R10-1.0 |
| PBS | Polarizing beamsplitter | Thorlabs | WPBS254-VIS |
| $\lambda_1/2$ | 488 nm zero-order half-wave plate | Thorlabs | WPHSM05-488 |
| $\lambda_2/2$ | 561 nm zero-order half-wave plate | Thorlabs | WPH10M-561 |
| $\lambda_3/2$ | 633 nm zero-order half-wave plate | Thorlabs | WPH10M-633 |
| $\lambda/4$ | 350 - 850 nm achromatic quarter-wave plate | Thorlabs | AQWP10M-580 |
| Ex | Quadband excitation filter | Chroma Technology | ZET405/488/561/640xv2 |
| Em1, Em2 | Quadband emission filter | Chroma Technology | ZET405/488/561/640mv2 |
| DM1,2,3 | Quadband dichromatic mirror | Chroma Technology | ZT405/488/561/640rpcv2 |
| DM4 | 550 nm short pass dichromatic mirror | Thorlabs | DMSP550R |
| DM5 | 605 nm long pass dichromatic mirror | Thorlabs | DMLP605R |
| Slit | Adjustable mechanical slit | Thorlabs | VA100 |
| Galvo1,2 | 5 mm aperture, VIS dielectric-coated galvanometric scanner | ScannerMAX | Saturn-5 |
| Camera | sCMOS camera | Teledyne Photometric | Kinetix |
| DMD | Digital micromirro device | ViALUX GmbH | V-7000 VIS |
| 488 nm Laser | 55mW 488nm diode laser | Lasertack GmbH | PD-01376 |
| 561 nm Laser | 200mW 561nm CW DPSS Laser | Oxxius | LCX-561L-200-CSB-PPA |
| 637 nm Laser | 6W 637 nm diode laser bar | Ushio America, Inc. | Red-HP-63x |

**Table S3.** List of experimental parameters.

| Dataset | GEVI | Imaging depth | Frame rate | Camera mode | Excitation intensity |
|---|---|---|---|---|---|
| Fig. 1(h,i) | somArchon | 100 µm | 500 Hz | Sensitivity | 2 $W/mm^2$ |
| Fig. 1(j,k) | Voltron2 | 150 µm | 800 Hz | Speed | 80 $mW/mm^2$ |
| Fig. 3 | Voltron2 | 160 $\pm$ 20 µm | 800 Hz | Speed | 40 $mW/mm^2$ |
| Fig. 4(a-c) | Voltron2 | 200 µm | 800 Hz | Speed | 80 $mW/mm^2$ |
| Fig. 4(d-f) | Voltron2 | 250 µm | 800 Hz | Speed | 120 $mW/mm^2$ |
| Fig. 4(g-i) | Voltron2 | 300 µm | 730 Hz | Sensitivity | 120 $mW/mm^2$ |
| Fig. 5 | Voltron2 | N/A | 800 Hz | Sensitivity | 80 $mW/mm^2$ |
| Fig. S12 | Voltron2 | 160 $\pm$ 20 µm | 800 Hz | Speed | 40 $mW/mm^2$ |
| Fig. S14,S14 | Voltron2 | 150 $\pm$ 20 µm | 800 Hz | Speed | 60 $mW/mm^2$ |
| Fig. S15 | somArchon | 100 µm | 775 Hz | Sensitivity | 2.5 $W/mm^2$ |
| Fig. S16 | somArchon | 80 µm | 800 Hz | Sensitivity | 3.5 $W/mm^2$ |
| Fig. S17 | somArchon | 80-100 µm | 800 Hz | Speed | 2 - 2.5 $W/mm^2$ |
| Fig. S19(a) | Voltron2 | 220 µm | 800 Hz | Speed | 80 $mW/mm^2$ |
| Fig. S19(b) | Voltron2 | 270 µm | 800 Hz | Sensitivity | 150 $mW/mm^2$ |
| Fig. S19(c) | Voltron2 | 300 µm | 800 Hz | Speed | 130 $mW/mm^2$ |
| Fig. S19(d) | Voltron2 | 300 µm | 800 Hz | Speed | 150 $mW/mm^2$ |
| Fig. S20 | Voltron2 | N/A | 800 Hz | Speed | 60 $mW/mm^2$ |
| Fig. S21 | Voltron2 | 150 µm | 1000 Hz | Speed | 80 $mW/mm^2$ |
| Fig. S22(a-e) | Voltron2 | 130 µm | 2000 Hz | Speed | 130 $mW/mm^2$ |
| Fig. S22(f-j) | somArchon | 80 µm | 4000 Hz | Speed | 14 $W/mm^2$ |
